# DNA methylation-based prediction of differentiation state at cell-type resolution reveals an age-associated epigenetic mesenchymal drift

**DOI:** 10.64898/2026.10.10.758162

**Authors:** Andrew E Teschendorff, Huige Tong, Xiaolong Guo

**Author notes:** Corresponding author: Andrew E. Teschendorff. Equal Contribution (Andrew E Teschendorff, Huige Tong).

## Abstract

Quantifying the differentiation state of cells is an important task in regenerative medicine, cellular reprogramming and in elucidating how differentiation hierarchies are altered in aging and disease. For DNA methylation data (DNAm) there is however an accute shortage of methods for quantifying differentiation state. Here we use machine learning to build EpiDiff, a predictor of differentiation that yields a continuous absolute differentiation index (DI) when applied to a DNAm sample. We extensively validate EpiDiff in single-cells, sorted cells and bulk tissue, as well as in *in-vitro* differentiation, organoid, reprogramming and cellular rejuvenation experiments. In mixed samples, EpiDiff measures the extent of differentiation across all cellular lineages. In sorted cell populations, we find that DI increases with age and, underpinning this increase, we discover that many cell-types undergo a developmental epigenetic drift that is most prominent towards the mesenchymal lineage. When applied to normal-tissues, EpiDiff is predictive of carcinogenic exposure and cancer risk. In summary, EpiDiff is a unique, general and versatile tool that can accurately quantify the global differentiation state of a DNAm sample with broad applicability to many disciplines.

---

DNA methylation (DNAm) plays a key role in early development, differentiation and is integral for tissue homeostasis ^1–4^. DNAm is also known to change with age ^5–10^, in normal tissues exposed to environmental risk factors ^11–14^, in preneoplastic lesions like dysplasia ^15^ and in diseases like cancer ^15–24^. These epigenetic changes may underpin changes in the differentiation state and hierarchy of tissues, predisposing tissues to disease ^25^. Thus, using DNAm to quantify the differentiation state of a sample could help improve our understanding of cellular aging and disease predisposition. For instance, a predictor of differentiation could be used in conjunction with a DNAm-clock ^26^, to assess how well a partial reprogramming protocol ^27–29^ rejuvenates cells without loss of cell-identity. A DNAm-based predictor of differentiation state could also help identify important age-related processes that underpin biological aging of cell-types ^26^. In the context of precancerous lesions ^16,30^, being able to infer differentiation state could help identify preneoplastic samples at higher cancer risk ^31,32^. Importantly, because DNAm can be measured with a high degree of accuracy ^33,34^, the inference of differentiation state from DNAm data could lead to a more accurate identification of disease risk states compared to noisier data-modalities like mRNA expression. Indeed, with Illumina DNAm beadarrays it is possible to reproducibly detect small 1-2% DNAm changes in relation to factors such as age ^9,35^ or smoking ^36^. Because of its relatively low cost, the overwhelming majority of DNAm data in the public domain has been generated with such beadarrays. Yet, despite this large-scale availability of DNAm data, there is no general DNAm-based method for quantifying differentiation-state.

Here we address this urgent gap in the epigenomics field, assembling a highly curated collection of DNAm datasets, all profiled with the same underlying technology, and subsequently using machine learning to build an accurate predictor of differentiation state called EpiDiff. We extensively validate EpiDiff in many independent DNAm datasets, representing different technologies, including single-nuclei DNAm data. Importantly, our predictor is robust to batch effects and yields a continuous differentiation index on an absolute scale, allowing inter dataset comparisons. We then illustrate the broad applicability of EpiDiff to cellular rejuvenation, aging and cancer risk prediction. One key finding is that many cell-types display age-associated drifts, reflecting DNAm changes that mimic those seen during differentiation of hESCs into fetal mesoderm, endoderm and ectoderm, with a mesenchymal drift being most prominent. Underpinning these drifts, we discover that CpGs implicated in early fetal differentiation represent age-associated hotspots of epigenetic dysregulation.

## Results

### EpiDiff: a predictor of differentiation state from DNAm data

To build a predictor of differentiation state we assembled a collection of Illumina DNAm datasets comprising 146 samples from three key differentiation states including pluripotency, multipotency (fetal tissue-types) and fully differentiated (adult tissue-types) (**Methods**). This collection included a total of 14 fetal and 14 adult tissue-types, of which 11 were matched (or approximately matched) (**Fig.1a, Methods**). We devised a CpG selection strategy to maximize the chances of identifying features that are truly associated with differentiation and not confounded by cell-division or batch effects (**Methods**). This strategy required, not only statistically significant differences in DNAm between differentiation states, but also that the effect sizes (absolute difference in average DNAm) are large enough to ensure that CpGs are tracking differentiation and not cell-division or batch effects (**Methods, Fig.1b, SI fig.S1**). Focusing on regulatory regions, we identified a total of 1298 differentiation-associated CpGs (‘diff-CpGs’) (**SI tables S1-S2**), of which the majority (882) were methylated in hESCs and became progressively hypomethylated with differentiation, with the rest (416) being unmethylated in hESCs and displaying hypermethylation with increased differentiation (**Fig.1c**). Our feature selection strategy ensured that diff-CpGs displayed significant DNAm differences between fetal-tissue and hESCs, as well as between adult tissue and hESCs, whilst also favoring CpGs that display progressive DNAm changes, i.e. with the fetal state displaying intermediate values between hESCs and adult tissue (**Fig.1d**). Gene Set Enrichment Analysis (GSEA) with two different methods (ebGSEA ^37^ and missMethyl ^38^) confirmed that the 416 hypermethylated CpGs were strongly enriched for developmental terms, including bivalent and PRC2 marked sites (**Methods, SI table S3-S4**). Interestingly though, the more numerous 882 hypomethylated CpGs were most strongly enriched for immune-response related terms including complement pathways, IL2-STAT5 signaling and inflammatory response (**SI table S3-S4**).

**Figure 1:**
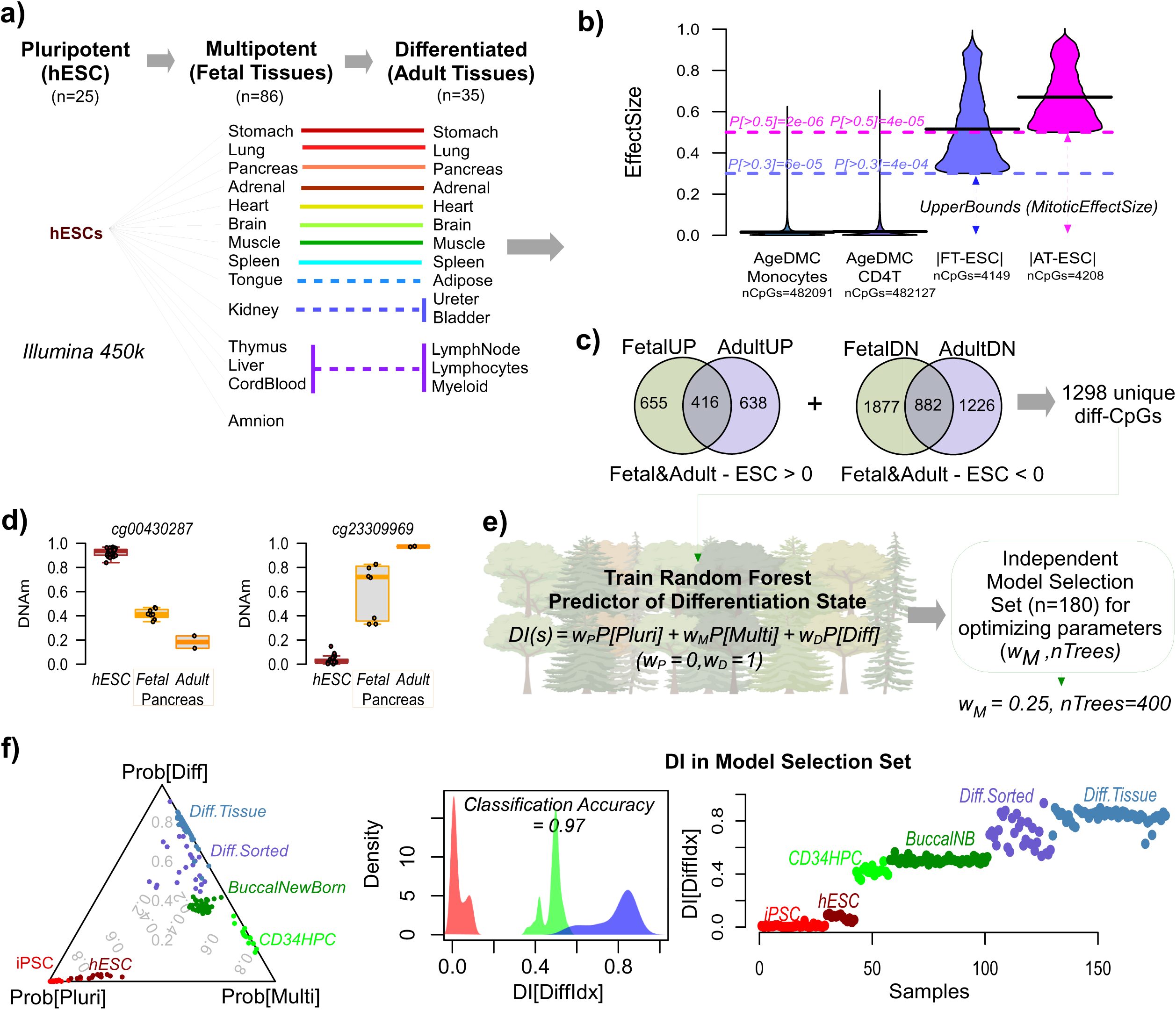
Construction and validation of EpiDiff. **a)** Diagram of pluripotent, fetal and adult tissue samples (mostly derived from the Stem-Cell Matrix Compendium (SMC2)) which we used for identifying CpGs implicated in differentiation. All DNAm data was processed uniformly with the same Illumina 450k technology. **b)** Violin plots illustrate our strategy for identifying CpGs where DNAm changes because of differentiation and not because of cell-division. Violin plots compare the effect size (DNAm-scale) of age-associated DMCs as derived from the Illumina 450k MESA DNAm dataset that profiled sorted monocytes (n=1199) and CD4T-cells (n=212), to the effect sizes of CpGs implicated in differentiation. In the case of age-DMCs, the effect size is the estimated regression slope multiplied by 80 years, to yield the typical change of DNAm during a human lifetime. Because cell-division is only one component that contributes to age-associated DNAm changes, this effect size distribution provides a conservative upper bound on how big DNAm changes can arise purely because of cell-division. In the case of the SCM2 data, effect sizes were computed as the mean change in DNAm between fetal tissues and hESCs, and between adult tissues and hESCs, as shown. Horizontal dashed lines indicate the thresholds chosen to select ‘differentiation-CpGs’. The probabilities within the panel indicate the fraction of 450k CpGs displaying an age effect size larger than these thresholds. **c)** Venn diagrams display the overlapping differentiation CpGs derived by comparing fetal tissue to hESCs with those derived comparing adult tissue to hESCs, stratified according to direction of DNAm change. **d)** Examples of 2 differentiation CpGs implicated in the differentiation of the pancreas. **e)** Diagram illustrating the training of a Random Forest classifier to predict differentiation state as a continuous differentiation index (DI) calculated from the probabilities of a sample belonging to pluripotent, multipotent or differentiated state. Two key parameters to optimize are the weighting scheme in the DI calculation and the number of trees. These parameters are optimized with an independent model selection set. **f)** Left: ternary plot of the predicted probabilities in the model selection set and for the optimal choice of parameters. Middle: density plot of the DI-values for samples in the model selection set, stratified by the known differentiation state. Right: Scatterplot of predicted DI-values in the model selection set.

Having identified a high confidence set of diff-CpGs, we next aimed to build a multi-class predictor of differentiation state, focusing on the aforementioned three differentiation states (pluripotency, multipotency and terminally differentiated). We reasoned that including pluripotent, fetal-tissue (as a surrogate for multipotency) and terminally differentiated adult samples would allow us to build a continuously valued differentiation index (DI) with values close to 0 representing samples/cells that are pluripotent and DI-values close to 1 representing terminally differentiated samples/cells. To build the DI-predictor we applied RandomForests (RF) ^39^ to the full training set using a range of different parameter settings, focusing on the selected 1298 diff-CpGs (**Methods, Fig.1e, SI fig.S2a**). In order to not overfit, we selected parameters that optimized performance across 5 independent DNAm datasets comprising a total of 180 samples (42 pluripotent: 29 iPSC samples from the SCM2 compendium ^40^ and 13 hESC samples from Butcher et al ^41^, 59 multipotent: 44 buccal swabs from newborns ^42^ and 15 CD34+ hematopoietic progenitor cells ^43^ and 79 differentiated: 28 sorted samples from 11 different adult tissue-types from the DNAm-atlas of Moss et al ^44^ and 51 samples from 9 adult tissue-types from Slieker et al ^45^) (**Methods, Fig.1e, SI fig.S2a**). Whilst a model selection set should ideally include more diverse multipotent samples, we wanted to use the few remaining multipotent datasets for fully blinded validation. We observed excellent validation performance of the RF-predictor in the model selection set of 180 samples (Classification Accuracy = 0.97, **Fig.1f**), which was also robust to the specific parameter values (**SI fig.S2b**), indicating strong generalization performance and little overfitting. To confirm that our DI-predictor is not confounded by cell-division we applied it to an independent EPIC DNAm dataset of 7 cell-lines, with each cell-line undergoing 30 to 60 population doublings (PDs) ^46^: predicted DI-values were stable and did not change appreciably or monotonically with PDs (**SI fig.S3a**). Moreover, when applied to the multi normal-adjacent tissue TCGA DNAm dataset, we observed that the DI displayed no correlation with the cumulative number of stem-cell divisions, nor with the tissue’s intrinsic stem-cell division rate ^47,48^ (**SI fig.S3b**), in stark contrast to a mitotic clock like stemTOC ^49^ (**SI fig.S3c**). Importantly, the DI index resolved differences in differentiation between samples within the same differentiation state: for instance, the DI-values for the CD34+ multipotent hematopoietic progenitor cells (HPC) were lower compared to those of newborn buccal swabs (**Fig.1f**), consistent with the latter samples only being enriched for multipotent cells. We call the final optimized RF-predictor, EpiDiff.

### Validation of EpiDiff in independent DNAm datasets

To demonstrate generalizability of EpiDiff we first applied it to iPSCs generated via reprogramming experiments (**Methods**). In all cases, the iPSCs displayed very low DI values and were correctly predicted to be pluripotent (**Fig.2a**). The original somatic cells were mostly categorized as differentiated but in some instances also as multipotent: for instance, foreskin fibroblast (FFib) samples from newborns were categorized as multipotent, consistent with these cells being more proliferative and potent than dermal fibroblasts from adults (**Fig.2a**). Adult dermal fibroblasts from Zellwegger-Syndrome-Disorder patients also displayed lower than expected DI-values, probably reflecting their disease state. To validate EpiDiff in sorted cells, we applied it to 7 sorted immune cell-types from cord-blood and 12 immune cell-types from adult blood, encompassing 220 samples derived from 5 independent DNAm datasets ^50,51^. As required, the DI of all adult immune cell-types were classified as differentiated and were higher than those of all cord-blood immune cell-types (**Fig.2b**). It is noteworthy that unlike cord-blood lymphocytes, which were classified as multipotent, monocytes and granulocytes from cord blood were classified as differentiated, consistent with innate immune cells from cord-blood being more mature than cord-blood lymphocytes ^50^. Further validating EpiDiff, adult naïve immune cell-types displayed lower differentiation levels than their corresponding memory counterparts (e.g. naïve B vs memory B, **Fig.2b, SI fig.S4a**).

**Figure 2:**
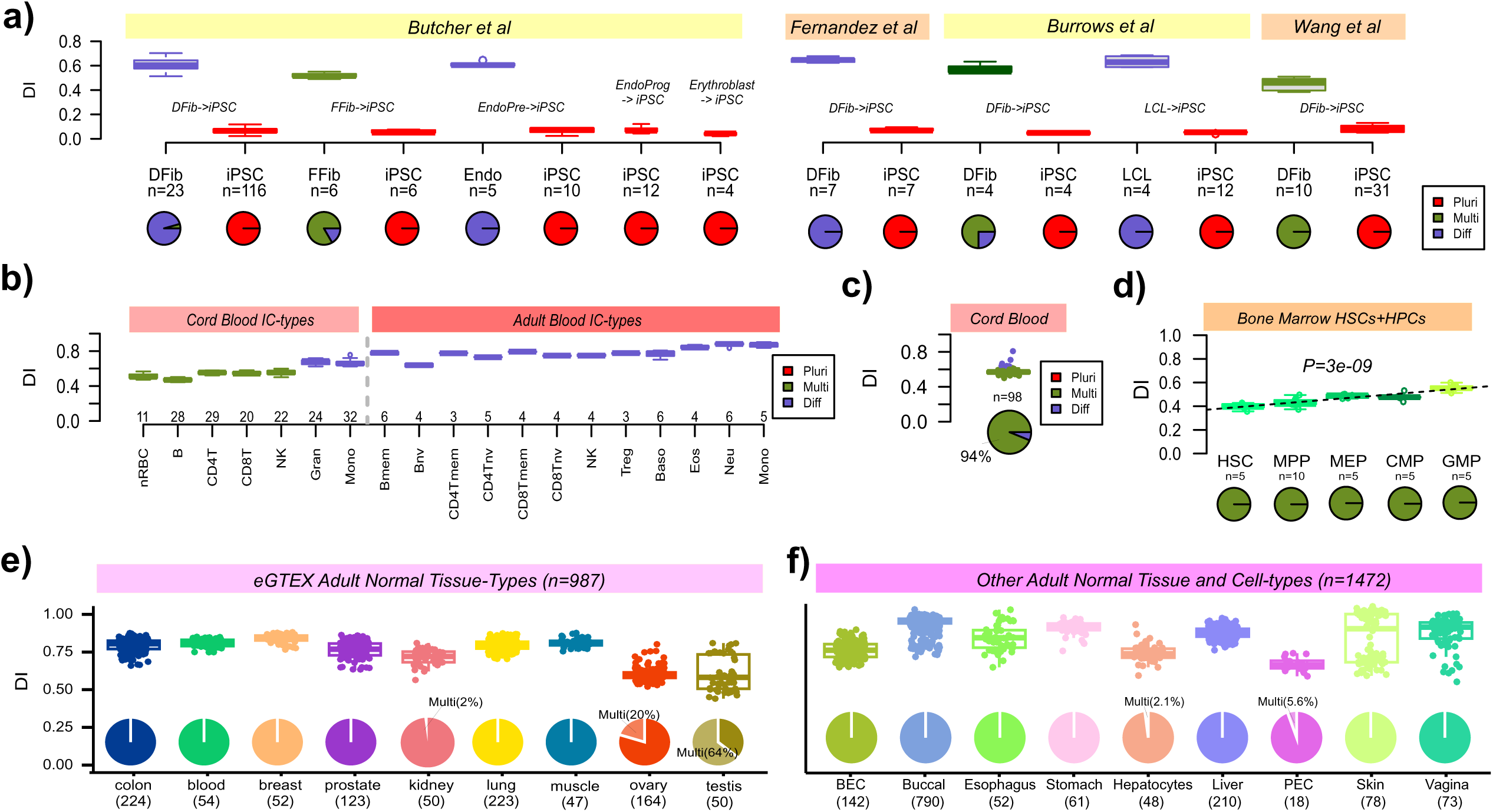
Independent validation of EpiDiff in Illumina DNAm data. **a)** Boxplots displaying the validation of EpiDiff (DI) in iPSC reprogramming DNAm data. Piecharts at the bottom show the distribution of predicted differentiation states. (DFib=dermal fibroblast, FFib=foreskin fibroblast from neonate, LCL=lymphoblast cell-line, iPSC=induced pluripotent stem-cell). **b)** Boxplots displaying the validation of EpiDiff (DI) in sorted cord and adult blood immune cell-types. Color of the boxes indicate the predicted differentiation state. **c)** Boxplot displaying the validation of EpiDiff (DI) in cord blood tissue samples, with piechart indicating the distribution of predicted differentiation states. **d)** As c) but for bone marrow derived hematopoietic stem and progenitor cell-types, with piecharts indicating the distribution of predicted differentiation states (all multipotent). HSC=hematopoietic stem cells, MPP=multipotent progenitors, MEP=megakaryocyte-erythrocyte progenitors, CMP=common myeloid progenitors, GMP=granulocyte myeloid progenitors. Linear regression fit and P-value are shown. **e-f)** Boxplots displaying the validation of EpiDiff (DI) in the adult normal tissue-types from eGTEX, as well as in an independently curated large collection of adult normal tissue-types. Piecharts show the distribution of predicted differentiation states. Number of samples per tissue-type is shown in brackets.

To further test the generalizability of EpiDiff, we applied it to a large series of 98 cord blood samples (Illumina 450k) ^52^. Their predicted DI-values were in line with those of the sorted cord-blood cell-types, with 94% of the samples being predicted as ‘multipotent’ (**Fig.2c**). We also applied EpiDiff to bone marrow derived hematopoietic stem and progenitor cells from healthy donors (n=30, **Methods**) ^43^. For all 30 samples, the predicted class was correctly predicted to be multipotent (**Fig.2d**). Importantly EpiDiff was sensitive enough to capture the subtle increase in differentiation from hematopoietic stem cells (HSC) to multipotent progenitors (MPP), and from MPPs to other more committed progenitor subtypes (**Fig.2d**).

To validate EpiDiff in adult tissue-types, we applied it to the large eGTEX DNAm EPIC dataset ^53^ profiling normal adult tissues, encompassing 9 tissue-types and a total of 987 samples. As required, the DI was high in effectively all tissue-types, with the overwhelming majority of samples assigned to the differentiated category (**Fig.2e**). Two exceptions were ovary and testis, which are known to contain a substantially higher fraction of more potent cells, thus consistent with their predicted reduced differentiation levels (**Fig.2e**). We further validated EpiDiff in other adult normal tissue-types, as well as in adult sorted samples representing bronchial epithelial cells (BECs), pulmonary endothelial cells (PECs) and hepatocytes (**Methods, Fig.2f**). Of note, although sorted cell-types like PECs and hepatocytes displayed lower DI-values, over 94% of samples were still correctly classified as fully differentiated (**Fig.2f**). For skin, we observed a bi-modal distribution (**Fig.2f**), with epidermal samples displaying lower DI-values than dermal ones (**SI fig.S4b**), in line with the known higher fraction of undifferentiated keratinocytes in the epidermis ^54^. Given that undifferentiated and differentiated keratinocytes in the epidermis fall along a unipotent lineage, this also suggests that EpiDiff captures differentiation gradients within unipotent lineages. To confirm this further, we applied EpiDiff to a multi-platform 450k & EPIC DNAm dataset profiling colon crypts (enriched for undifferentiated unipotent epithelial stem-cells) and colonic mucosa (enriched for differentiated progeny), revealing significantly lower DI values in the crypts (**SI fig.S4c**). Importantly, the DI-values of samples generated with 450k and EPIC platforms were statistically indistinguishable, confirming that EpiDiff is robust to platform differences (**SI fig.S4c**). In summary, these data clearly demonstrate the ability of EpiDiff to predict the differentiation state of reprogrammed cells, sorted cells and bulk-tissue, and across two different Illumina DNAm technologies (450k & EPIC).

### Validation of EpiDiff in *in-vitro* differentiation experiments

To assess if EpiDiff can track differentiation in *in-vitro* differentiation experiments, we applied EpiDiff to an experiment (EPIC arrays) that used 4 hepatocyte-specific TFs (4F) to differentiate umbilical cord blood derived mesenchymal stem-cells (MSC) to hepatocyte-like cells ^55^ (**Methods, Fig.3a**). Reassuringly, the DI value of the starting MSC population was classified as multipotent, whilst differentiation into hepatocyte-like cells led to an increase in DI, with a corresponding switch in probability class assignments indicating a significantly more differentiated state (**Fig.3a**). Importantly, when the MSCs were treated with a non-differentiation inducing factor, the DI value remained low (**Fig.3a**). We also applied EpiDiff to an *in-vitro* human brain organoid model of cortex development where human cortical spheroids were cultured from human induced pluripotent cell-lines derived from 4 donors encompassing 13 timepoints (total of 50 samples profiled with EPIC DNAm arrays) ^56^ (**Methods**). For each cell-line, EpiDiff’s DI increased with differentiation timepoint (**Fig.3b**). Of note, the final attained DI values were still lower compared to those of adult post-mortem cortex tissue, consistent with a postnatal pre-adult maturation stage (**Fig.3b**). Overall, these data indicate that EpiDiff has the sensitivity to track differentiation changes *in-vitro*.

**Figure 3:**
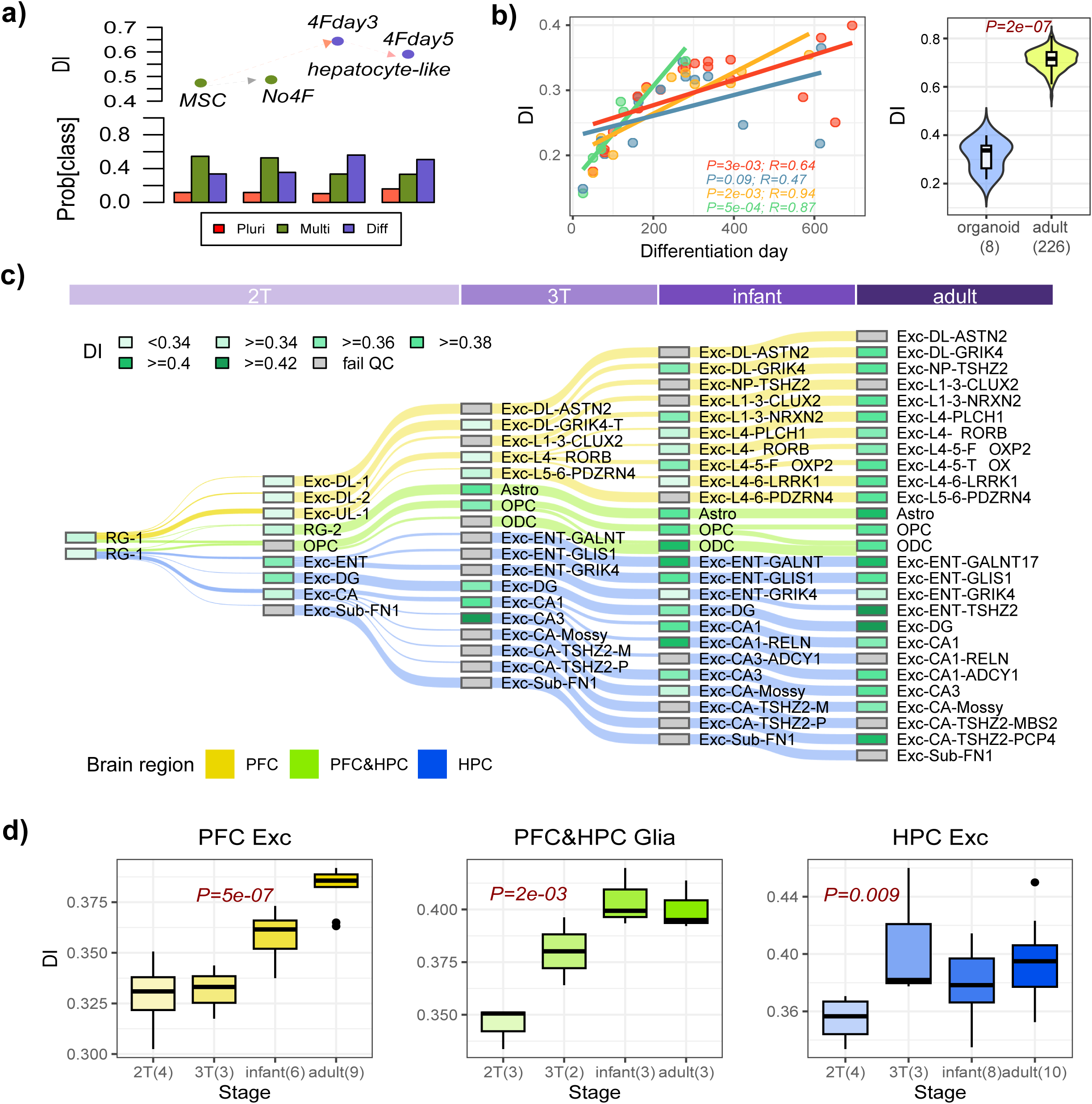
Validation of EpiDiff in in-vitro differentiation and snmC3-Seq3 data. **a)** Predicted changes of DI in an experiment where mesenchymal stem cells (MSCs) were differentiated using 4 differentiation factors (4F) into hepatocytes (4Fday3 and 4Fday5). “No4F” means no factors were applied which serves as negative control. Barplots compare the predicted differentiation state class assignments for the 4 samples. **b)** Predicted DI values against differentiation day for an in-vitro organoid differentiation experiment of human induced PSCs into human cortex (up until postnatal stage) for 4 different donors (labeled by colors). Regression fits for each donor are shown, as well as the R-values and correlation test P-values. Donor IDs: 1205 (n=11), 0307 (n=7), 2242 (n=14), 8858 (n=19). Violin plots to the right compare the DI values of the organoid experiment (last two timepoints for each of 4 donors, making 8 samples) to 226 adult pre-frontal cortex samples from Jaffe et al. P-value is from a one-tailed Wilcoxon rank sum test. **c)** Alluvial/Sankey diagram of the developmental lineage hierarchy of approximately 56 brain cell-types during development of the human brain, including prefrontal cortex (HPC) and hippocampus (HPC) regions. Developmental stages cover mid-gestation (2^nd^ trimester-2T), late gestation (3^rd^ trimester-3T), infants and adults. Each cell-type is labeled with a color according to their predicted DI-value, as indicated. RG=radial glia, Astro=astrocyte, OPC=oligo-progenitor-cell, ODC=oligodendrocyte, Exc=excitatory neuron. For other cell-type annotations, see Methods. **d)** Boxplots displaying the change in DI-values across developmental stages for excitatory neuron lineage in HPC and PFC, as well as that of the glia cells shared by both HPC and PFC. P-values for PFC-Exc and glia lineages derive from a linear regression test, whilst for HPC-Exc, it is derived from a one-tailed Wilcoxon-test comparing 2T stage to the rest.

### Validation of EpiDiff on single-cell DNAm (snm3C-Seq3) data

Next, we asked if EpiDiff is applicable to single-cell DNAm data. To this end, we obtained single-nucleus methyl-3C (Chromatin Conformation Capture) sequencing (snm3C-Seq3) data of 53,000 single nuclei from the developing human brain, comprising prefrontal cortex (PFC) and hippocampus (HPC) samples from mid-gestation (2T), late-gestation (3T), infants and adults ^57^. Due to the sparseness of snm3C-Seq3 data, we analyzed normalized pseudo-bulk aggregates for approximately 10 brain cell-types using the annotation provided by Heffel et al ^57^. SVD analysis of the pseudo-bulk data confirmed segregation of brain cell-types according to lineage and developmental stage (**SI fig.S5a**). Of the 1298 EpiDiff CpGs, an average of 89% CpGs could be mapped to each pseudo-bulk profile passing QC (**SI fig.S5b, Methods**). Next, we arranged the brain cell-types along the developmental lineage hierarchy proposed by Heffel et al ^57^, overlaying onto each sample the predicted DI value from EpiDiff (**Fig.3c**). Despite the missing CpGs and different technology, EpiDiff correctly predicted increased DI-values along the various lineage trajectories (**Fig.3c**). To test the statistical significance of this, we applied linear regression and non-parametric testing to the differentiation lineage of excitatory neurons from the prefrontal cortex (PFC) and hippocampus (HPC) regions, revealing in both cases a statistically significant increase (**Fig.3d**). This pattern was also validated for glia’s differentiation lineage trajectory, present in both PFC and HPC (**Fig.3d**), as well as for the inhibitory neuronal lineage **(SI fig.S6**). Overall, these results demonstrate that EpiDiff is applicable to snm3C-Seq3 data.

### Invariance of DI following transient reprogramming and cellular rejuvenation

An important test and application of EpiDiff is to cellular rejuvenation experiments using transient reprogramming ^29,58^. We first applied EpiDiff to an EPIC DNAm dataset generated using a transient reprogramming protocol termed OSKLMN, according to which somatic dermal fibroblasts and endothelial cells were treated with a cocktail of mRNAs representing OCT4, SOX2, KLF4, LIN28, c-MYC and NANOG for 4 consecutive days (in contrast to full reprogramming which would involve treatment for 12-15 days) ^58^. Whilst Horvath’s pan-tissue clock DNAmAge was reduced in the treated ‘rejuvenated’ cells (**Fig.4a**), the DI remained invariant (**Fig.4b**), consistent with a protocol that rejuvenates cells without a major loss of cell-identity. Next, we applied EpiDiff to a different reprogramming experiment where young fibroblasts were treated with OSKM+doxycycline for a maximum of 17 days (maturation phase), subsequently undergoing further treatment to a fully reprogrammed state ^29^. In a separate maturation phase transient reprogramming (MPTR) experiment, fibroblasts from old donors were also treated with OSKM+doxycycline for a maximum of 17 days till maturation phase, then flow-sorted according to SSEA4 (a marker of successful reprogramming) and subsequently grown without doxycycline, a protocol hypothesized to rejuvenate cells ^29^. In the first full reprogramming protocol, DNAmAge, as measured with Horvath’s skin&blood clock^59^ decreased, whilst the DI in the maturation phase only remained marginally lower than that of the young fibroblasts (**Fig.4c**). In the case of the MPTR experiment, DNAmAge displayed a marginal reduction after doxycycline removal (**Fig.4d**). Importantly, after doxycycline removal, the DI displayed high stability and remained relatively close in value to the one displayed by old fibroblasts (**Fig.4d**). Overall, these data suggest that EpiDiff predicts preservation of cell-identity upon transient reprogramming, although in the case of MPTR, ‘rejuvenated’ fibroblasts displayed a lower DI than the original fibroblasts.

**Figure 4:**
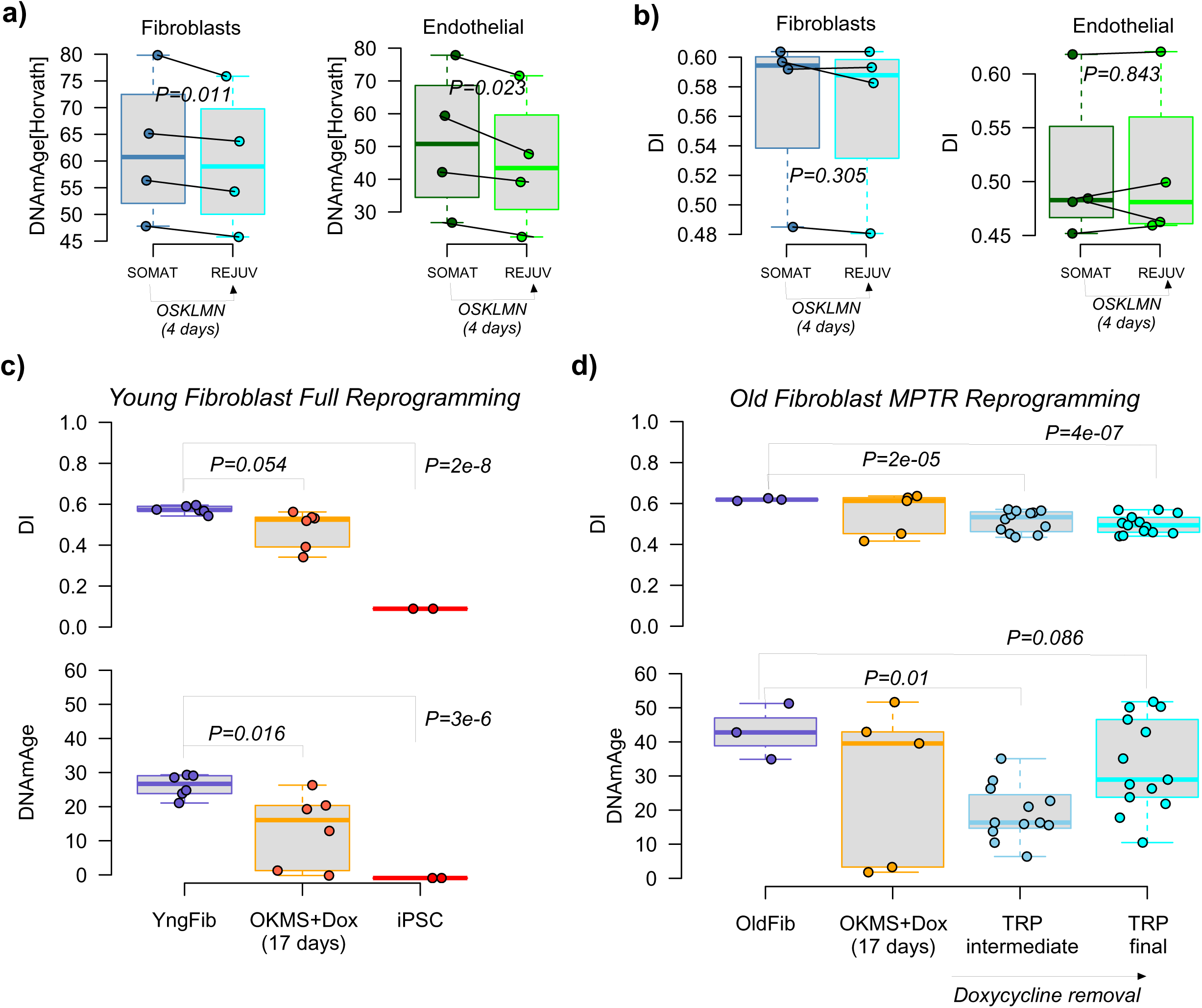
Validation of EpiDiff in cellular rejuvenation. **a)** Boxplots compare the predicted Horvath Clock DNAm-Ages of somatic cells (fibroblasts and endothelial cells) and their rejuvenated counterparts. P-value is from a one-tailed paired t-test. **b)** As a), but for the DI index with P-values derived from a two-tailed paired t-test. **c)** Boxplots of DI-values (top) and Horvath’s Skin&Blood-clock DNAmAge (bottom) in a full reprogramming experiment into iPSCs. P-values for DI and DNAmAge were derived using two-tailed and one-tailed t-tests, respectively. **d)** Top: Boxplots of DI-values in a maturation phase transient reprogramming (MPTR) of old fibroblasts (OldFib) into transiently reprogrammed fibroblast intermediates (TRPI) and transiently reprogrammed fibroblasts (TRP). The OKMS samples refer to application of the Yamanaka factors for 17 days. P-values are from a two-tailed t-test. Bottom: As top but for Horvath’s Skin&Blood DNAmAge. P-values are from a one-tailed t-test.

### EpiDiff measures global differentiation

Next, we wanted to assess the impact of cell-type heterogeneity on DI estimates obtained in bulk-tissue. For instance, sorted differentiated cells displayed a marginally lower DI than unsorted (mixed) differentiated samples (**Fig.1f**), suggesting that mixing together differentiated cell-types may increase the DI. To explore this, we collected DNAm datasets representing sorted immune ^60,61^ and brain cell-types ^62^, subsequently generating in-silico mixtures (**Methods**). In the case of mature immune cell-types, we mixed together the 15 possible pairwise combinations of 6 cell-types (B-cells, NK-cells, CD4+ and CD8+ T-cells, neutrophils and monocytes) (**Methods**). In each case, the mixed sample displayed a significantly higher DI than the original samples used in the mixture (**Fig.5a**). A similar pattern was observed when mixing together 3 brain cell-types, either with data generated with the Illumina platform (**Fig.5b, SI fig.S7a**), or with single-cell DNAm data (snmC-Seq2) ^63^ (**SI fig.S7b**). In the latter case, given the high resolution (single-cell) nature of snmC-Seq2 data, which allowed definition of 9 cell-types, we observed how DI increased in line with the number of cell-types in the mixture, with evidence of saturation when 7 or more cell-types were mixed together (**SI fig.S7b**). However, when we mixed together immature naïve immune cell-types (e.g. naïve CD4T-cells), which display lower DI-values than their memory counterparts (**SI fig.S4a**), with mature cell-types (e.g. monocytes), the mixed samples displayed intermediate DI-values (**SI fig.S7c**). Thus, we can reconcile these observations, concluding that EpiDiff is an aggregate measure of global differentiation, i.e. when applied to a mixed sample it not only reflects the extent of differentiation within a specific lineage, but also the aggregate extent of differentiation across all other lineages (cell-types).

**Figure 5:**
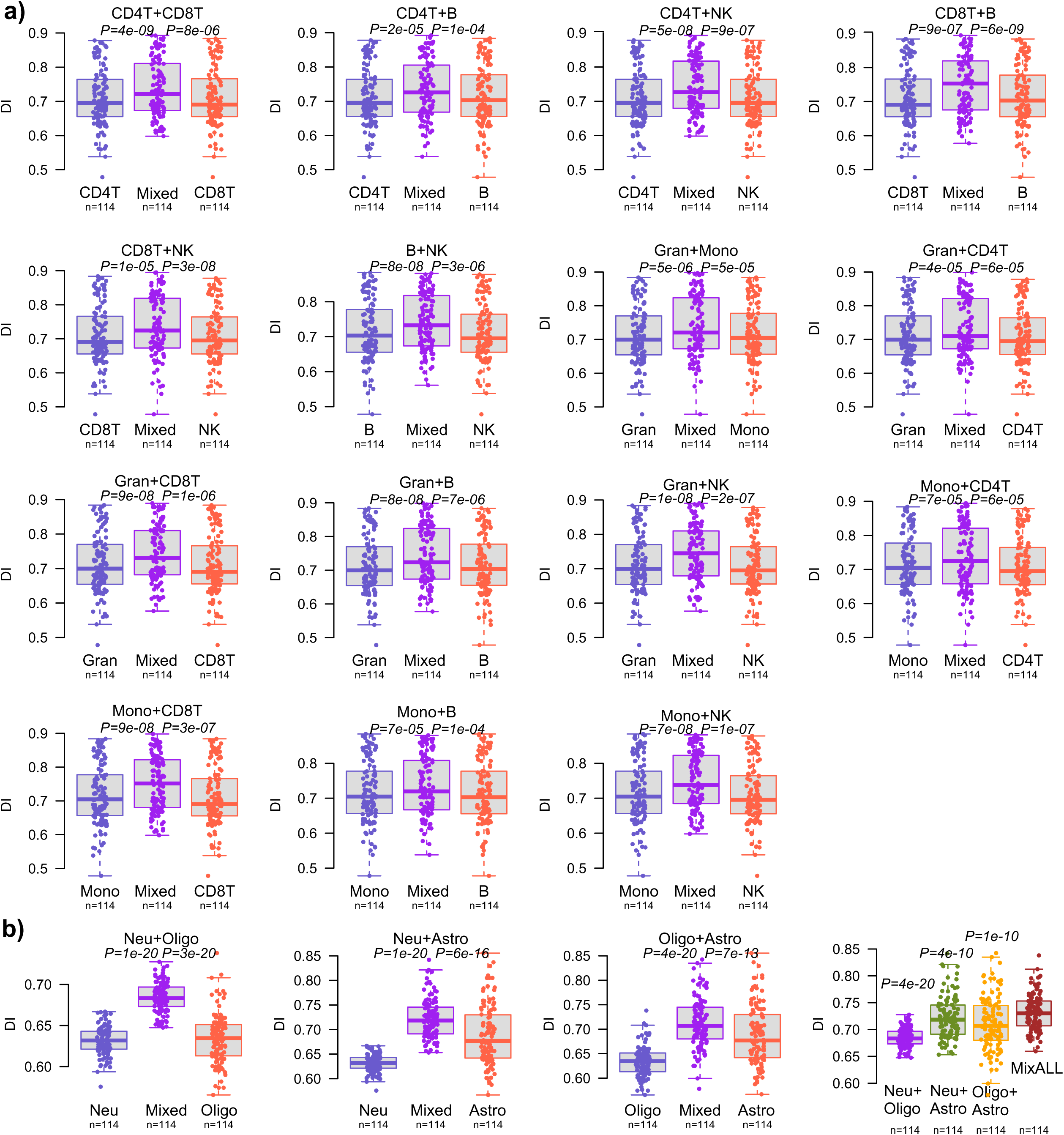
EpiDiff is a measure of global differentiation. **a)** Each panel compares the distribution of DI-values of a pair of immune cell-types to the mixture (“Mixed”) generated by mixing together the two sorted cell-samples from the same donor. Number of donors with matched numbers of sorted cell-type samples is given below each boxplot. Which cell-types are mixed is indicated above each panel. There are 15 panels in total for each pairwise combination of 6 immune cell-types (CD4+ and CD8+ T-cells, B-cells, NK-cells, Granulocytes and Monocytes). P-values compare the DI-values of the Mixed samples to those obtained in the sorted immune cell-types within each panel, and are derived using a paired one-tailed Wilcoxon rank sum test. DNAm EPIC data is from Bell et al. **b)** As a), but for DNAm data of sorted neurons, oligodendrocytes and astrocytes from Walker et al. The last panel compares the DI-values of mixtures of all 3 cell-types to the mixtures of 2 cell-types. P-values derived using a paired one-tailed Wilcoxon rank sum test.

### DI increases with age in sorted cells

Next, we asked if the differentiation index of cells changes with age. To this end, we applied EpiDiff to DNAm datasets profiling sorted cells from a wide age range. We first considered an EPIC DNAm dataset profiling 55 cultured hepatocyte cells, derived from adult donors aged 20 to 70 years, but also including 8 children under the age of 10. Remarkably, the DI was strongly associated with age (adjusted for sex, linear P=1e-11), with the young hepatocytes from children displaying significantly lower values compared to adults (**Fig.6a**). This indicates that hepatocytes from young children are far less differentiated and hence that they may retain higher potency, consistent with previous reports ^64^. Of note, the association with age remained significant when only considering adults (linear P=0.01 adjusted for sex). We next applied EpiDiff to a cohort of bronchial epithelial cells (BECs) ^65^, revealing an increased correlation with age, which remained significant after adjustment for asthma, sex and ethnicity (**Fig.6a**). When applied to sorted naïve CD4+ T-cells, monocytes and neutrophils from BLUEPRINT ^61^, EpiDiff revealed a strong association of DI with age in naïve CD4+ T-cells (**Fig.6a**). This increase would be consistent with a gradual increase in a more mature cell subpopulation, which however would still express the naïve CD4+ T-cell marker used for sorting. Confirming this hypothesis, we estimated the purity of the naïve CD4+ T-cells with our EpiDISH algorithm ^66^, finding a strong anti-correlation of DI with purity, whilst simultaneously also displaying a strong positive correlation with the estimated mature CD4+ T-cell fraction (**SI fig.S8a**). In contrast, myeloid cells are already relatively mature at birth ^50,67^, which may explain the relative constancy of DI with age in these cell-types (**SI fig.S8b**). Finally, we applied EpiDiff to one of the largest Illumina DNAm datasets representing 4 sorted brain cell-types (Neurons, Oligodendrocytes, Microglia and Astrocytes) ^62^. Interestingly, this revealed an increased DI with age in all glia subpopulations (**Fig.6a**), but not in neurons (**SI fig.S8c**). The non-significant pattern for neurons was also seen in two additional, albeit smaller, cohorts (**SI fig.S8c**, **Methods**). Of note, all these patterns were largely recapitulated when we applied EpiDiff to single-nucleus DNA methylation sequencing data (snmCT-seq) from the prefrontal cortex ^63^, a study which profiled a total of 777 astrocytes, 314 microglia, 1157 oligodendrocytes and 52531 neurons (including 34587 excitatory neurons and 17944 inhibitory neurons) derived from old (70-74 years old) and young (23-30 years old) donors (**Methods, SI fig.S9**). Overall, these data suggest that many cell-types display increases of DI with age with some exceptions like neurons and myeloid cells.

**Figure 6:**
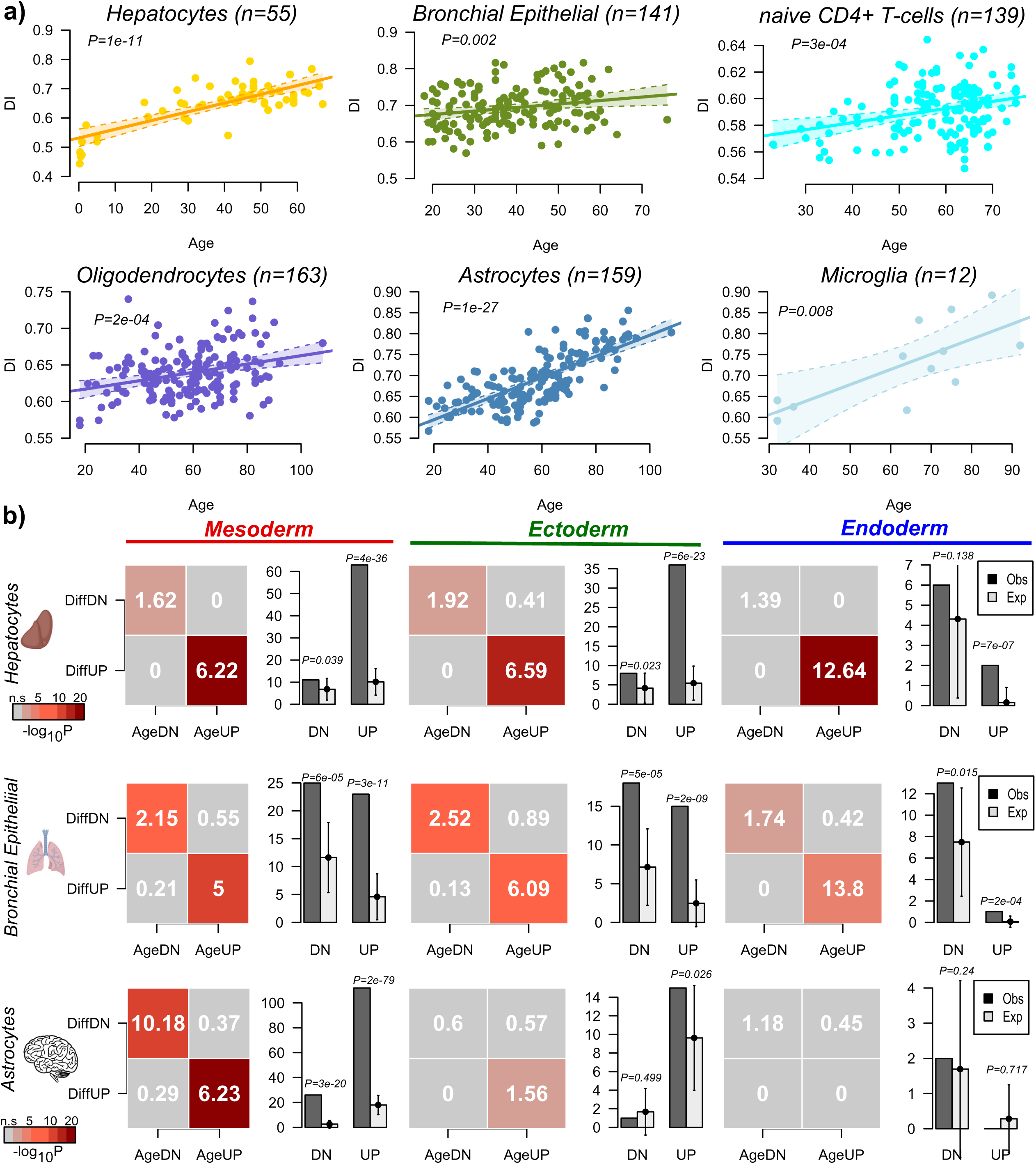
DI increases with age in sorted cell populations underpinned mostly by a mesenchymal drift. **a)** Application of EpiDiff to 6 Illumina EPIC/450k DNAm datasets of sorted cell populations, displaying the predicted DI-values against chronological age. Best linear regression fit adjusted for sex, 95% confidence interval envelope and P-value are shown. Number of sorted samples is given above each plot. **b)** For each germ-layer (mesoderm, ectoderm, endoderm) we display a heatmap of observed to expected ratios, one for each of four combinations of how germ-layer specific CpGs change with differentiation and with age in sorted hepatocytes (n=55), bronchial epithelial cells (n=141) and sorted astrocytes (n=159). Significance for calling significant age-DMCs in each dataset was determined at FDR<0.05. DiffUP/DiffDN: higher/lower methylation in fetal germ-layer compared to hESCs (derived from EpiDiff training set); AgeUP/AgeDN: age-associated hyper/hypo-methylation (derived from each sorted cell-type dataset). The color in the heatmap indicates statistical significance level, as shown. Barplot on the right compares the observed (black) to expected (lightgrey) number of CpGs that exhibit hypermethylation with both differentiation and age (UP), or hypomethylation with both differentiation and age (DN). For the expected number, we also display the 95% confidence interval.

### Pervasive age-associated mesenchymal drift underpins increase in global differentiation

In order to understand why the DI increases with age in sorted cells, we posited that this could reflect underlying cell-type heterogeneity driven by an age-associated developmental DNAm drift, which would include the germ-layer specific CpGs implicated in fetal differentiation. To explore this, we first assessed how many of the previously identified 416 hypermethylated and 882 hypomethylated differentiation-CpGs are specific to each germ-layer (**Methods**). We identified 247 mesoderm specific, 139 ectoderm specific and 70 endoderm specific differentiation CpGs (**SI tables S5-S6**). We then asked if and how these sites change with age in each of the sorted DNAm datasets analyzed earlier (**Methods**). This revealed a striking tendency for these CpGs to display age-related DNAm changes that are directionally aligned with how they change during differentiation (**Fig.6b, SI fig.S10**). Remarkably, in most cases, the observed number of hyper and hypomethylated germ-layer specific CpGs displaying respective age-associated hypermethylation or hypomethylation, was significantly higher than expected by random chance (**Fig.6b, SI fig.S10**), highlighting how these CpGs are hotspots of age-associated epigenetic dysregulation. Although this pattern was particularly pronounced and consistent for mesoderm-specifying CpGs, in many cases, the pattern of change was similar for different germ-layers. For instance, for liver hepatocytes and bronchial epithelial cells, the pattern was seen for diff-CpGs from all 3 layers (**Fig.6b**). On the other hand, for astrocytes the pattern was only seen for mesoderm-CpGs. Even in neurons, for which the DI did not clearly increase with age, meso- and-ectoderm specific CpGs were more likely to change with age than randomly selected CpGs (**SI fig.S10**). Findings were robust to the choice of significance threshold when calling significant age-associated changes (**SI fig.S11**). The same alignment of differentiation and age was also seen if we restricted to CpGs that are only implicated in the differentiation of fetal to adult tissue, i.e. without demonstrable DNAm changes between hESCs and fetal germ-layers (**Methods, SI fig.S12**). Overall, these data support the view that a significant fraction of the CpGs in EpiDiff display preferential age-associated DNAm changes and that they do so in a way that is directionally aligned with the changes during early differentiation, explaining why the DI increases with age.

### DI anti-correlates with mitotic age in cancer and smoking-exposed normal tissue

We next explored DI changes between normal and cancer tissue. Unlike mitotic age, which is universally increased in cancer compared to normal tissue ^49^, EpiDiff’s DI displayed significant decreases in 11 of 15 TCGA cancer-types (**SI fig.S13**). This is consistent with the reduction in differentiation of cancer cells compared to their normal counterparts. Importantly, when correlating stemTOC’s mitotic age to the DI using cancer samples only, we observed that for most cancer-types there was a significant anticorrelation, in line with more proliferative tumors being less differentiated (**Fig.7a**). In an independent Illumina 450k breast cancer DNAm dataset ^68,69^, we observed a similar decrease of DI in cancer compared to normal-tissue, a strong anticorrelation between mitotic age and DI across tumors, as well as a significant decrease of DI with histological grade (**Fig.7b**). Thus, these data demonstrate that a tumor’s DI and mitotic age are generally anticorrelated.

**Figure 7:**
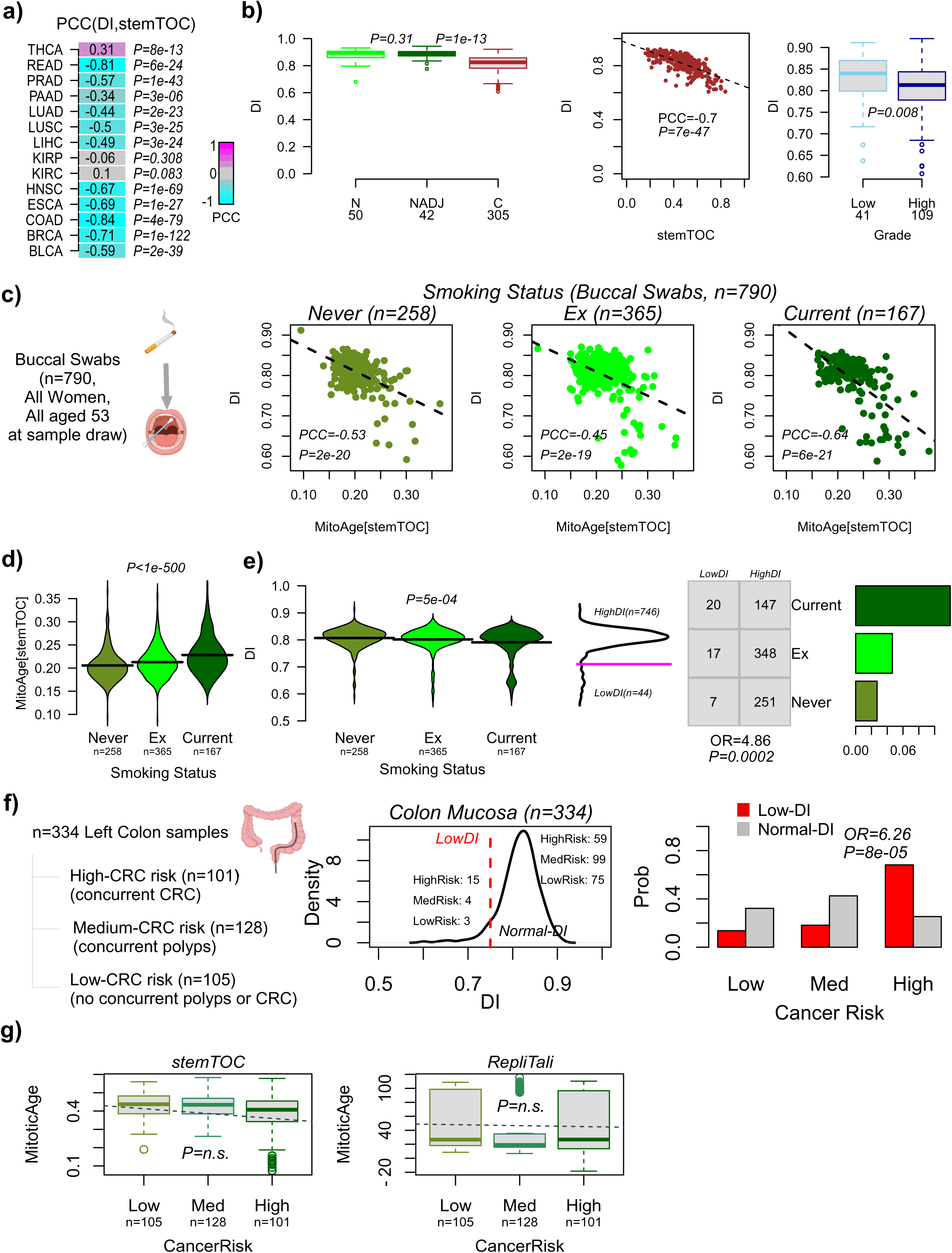
Lower DI marks increased mitotic age and cancer risk. **a)** Pearson Correlation Coefficients (PCC) between DI and stemTOC’s mitotic age as evaluated across TCGA tumor samples, with PCCs being displayed for each tumor-type. P-values derive from a correlation-test**. b)** DI-values against disease status for an Illumina 450k DNAm data of 50 normal breast samples (N), 42 normal-adjacent breast samples (NADJ) and 305 breast tumors (C). P-values are from a one-tailed Wilcoxon rank sum test, comparing N to NADJ and NADJ to C. Scatterplot of DI vs stemTOC’s mitotic age across all 305 breast tumor samples, with PCC and correlation test P-value. Boxplots of DI-values comparing low to high grade tumors. P-value is from a one-tailed Wilcoxon test. **c)** Scatterplots of DI-values vs mitotic-age (as estimated using stemTOC) in a buccal swab dataset of 790 samples from women all aged 53 at sample draw, with samples stratified by smoking-status (never, ex-smokers and current smokers). Pearson correlation coefficient (PCC) and P-values are given for each stratum. **d)** Violin plots of stemTOC’s mitotic-age vs smoking-status, with number of samples in each group given below plot. P-value derives from a linear regression of mitotic age vs smoking status (0=never, 1=ex, 2=current). **e)** Violin plots as in d) but for the DI-index. Density plot to the right of violin plots highlights the existence of two peaks, with the magenta line defining a threshold to divide samples up into high and low DI categories. Table displays how samples distribute according to smoking status and DI-state. Fisher test Odds Ratio and P-value are given. Barplot compares fraction of low DI samples stratified by smoking status, these differences being significant according to the Fisher-test of the table. Lower DI predicts colon cancer risk. **f)** Application of EpiDiff to an Illumina DNAm dataset of 334 colon mucosa samples, with individuals stratified into low, medium and high colorectal cancer (CRC) risk groups. Density plot displays the distribution of DI-values over all 334 samples. Deviations from Gaussianity as indicated by the threshold DI=0.75 (red-dashed line) classifies samples into lowDI and ‘normal DI’ groups. Number of samples in each risk group is indicated within the figure panel. Barplot to the right compares the probabilities (y-axis) of each risk group having a low-DI or normal-DI sample. Odds Ratio and associated P-value are derived from a one-tailed Fisher test comparing the high risk to the combined medium+low groups. **g)** Boxplots of mitotic ages (stemTOC and RepliTali) vs cancer risk group. P-values were estimated using a one-tailed regression test and were in both cases P>0.5, i.e. not significant (n.s.). P-values did not change appreciably when adjusting for age and sex. We used a one-tailed test because mitotic age should be higher in the high cancer risk group.

To see if this anti-correlation is also present in normal tissues exposed to a cancer risk factor, we applied EpiDiff and stemTOC to a large buccal swab DNAm dataset of 790 samples from women all aged 53 at sample collection and with highly variable smoking histories ^12^. We observed that DI and stemTOC’s mitotic age were strongly anti-correlated, specially in the group of current smokers (PCC=-0.64, P<1e-20, **Fig.7c**). Whilst mitotic age displayed a clear gradual increase from never-smokers to ex-and-current smokers (**Fig.7d**), the DI displayed a corresponding decrease, driven by outliers with low DI-values in the current smoker group (**Fig.7e**). To test the significance of this further, we thus applied a non-parametric one-tailed Fisher-test comparing the number of never and current smokers among low and high DI samples, which confirmed that current smokers displayed a significantly higher fraction of lower DI samples (OR=4.86, P=0.0002, **Fig.7e**). Thus, these results are consistent with buccal swabs exposed to cigarette smoke undergoing more rapid cellular turnover driven by amplification of a less differentiated progenitor pool. Overall, these data indicate that in the context of tumors and exposed tissues, differentiation and cell-proliferation are anti-correlated.

### DI in normal colonic mucosa correlates with colorectal cancer risk

We also applied EpiDiff to an Illumina DNAm dataset (n=334) of normal colonic mucosa samples, all drawn from the left colon, and with the 334 individuals stratified by low (n=105), medium (n=128) and high cancer-risk (n=101), according to absence or presence of concurrent adenomatous polyps and colorectal cancer (**Methods**) ^70^. Interestingly, the distribution of DI-values across all 334 samples was approximately Gaussian, except for a number of samples displaying lower DI-values **(Fig.7f**), reminiscent of the pattern seen in the buccal swabs from smokers. Strikingly, of the 22 samples displaying lowest DI-values, 15 (i.e. 68%) were high-risk individuals, whereas among the rest of samples, the fraction of high-risk individuals was only 25%, representing a highly significant shift in cancer risk (Odds Ratio= 6.3, Fisher test P=8e-5, **Fig.7f**). In contrast, application of two state-of-the-art mitotic clocks (stemTOC ^49^ and RepliTali ^46^) did not reveal the expected increased mitotic age with cancer-risk status, even after adjusting for age and sex (**Fig.7g**). Thus, these findings underscore the potential of EpiDiff to identify, from normal colon tissue, individuals at higher colorectal cancer risk.

### DI of pluripotent samples correlates inversely with their differentiation capacity

Finally, to explore the practical value of EpiDiff for regenerative medicine, we asked if EpiDiff can discriminate iPSCs according to their differentiation capacity (DC). To this end, we analyzed the DNAm data from Butcher et al who characterized hESC and iPSC samples in terms of their capacity to differentiate into the endodermal lineage ^41^. We found that the DI was significantly higher for those pluripotent samples deemed to be of lower differentiation capacity (**SI fig.S14a**). Of note, EpiDiff displayed more robust associations with DC than Horvath’s pan-tissue clock ^35^ (**SI fig.S14b**). To find out why EpiDiff is able to predict DC, we posited that those EpiDiff CpGs that become hypomethylated specifically during endoderm commitment would display lower DNAm in the pluripotent samples with higher endodermal differentiation capacity, which would be consistent with these pluripotent samples being inherently primed or skewed for endoderm differentiation. Hence, we computed t-statistics of differential DNAm for all the previously identified germ layer specific diff-CpGs (**SI table S5-S6, SI fig.S14c**) in the Butcher et al dataset, comparing all high DC to all low DC samples. Consistent with our hypothesis, the endodermal-specific CpGs displayed a strong skew towards lower DNAm levels in the high endoderm-DC samples, whereas this pattern was much weaker or non-significant for mesodermal and ectodermal CpGs (**SI fig.S14c**). We confirmed this pattern using either strict Bonferroni or relaxed nominal P<0.05 thresholds when calling differentially methylation: only endodermal CpGs displayed a clear skew towards lower DNAm levels in high endoderm-DC samples, whereas more mesodermal and ectodermal CpGs displayed hypermethylation (**SI fig.S14d**). This latter observation explains why the DI index, which is a global measure of differentiation (see next section) can be lower for high-endoderm DC samples whilst simultaneously displaying a skew or priming towards the endoderm lineage. In summary, these data demonstrate how EpiDiff has the resolution to distinguish pluripotent samples according to their DC.

## Discussion

Addressing a major gap in the epigenomics field, we have here successfully built a predictor of differentiation (EpiDiff) that is applicable to DNAm data generated with diverse technologies, including not only Illumina beadarrays but also snmCT-Seq and snm3C-Seq3. We have extensively validated this predictor across many independent DNAm datasets, comprising sorted cells, bulk-tissue, *in-vitro* differentiation and reprogramming experiments. As shown, EpiDiff is a versatile tool with broad applicability: for instance, in combination with an epigenetic clock, it can be used to assess how well an epigenetic reprogramming protocol rejuvenates cells without compromising cell-identity, with EpiDiff helping to quantify the degree of cell-identity preservation. EpiDiff was also able to predict differentiation capacity of iPSCs and to explain the variations in endoderm-differentiation capacity mechanistically in terms of the underlying DNAm patterns at endoderm-specific CpGs, a critical insight not found in previous studies using standard methods ^41^. We have also seen how EpiDiff was able to detect subtle changes in differentiation within buccal swabs of smokers or in the normal colonic mucosa of colorectal cancer patients. These shifts in differentiation state within normal tissues are consistent with increased cell proliferation of an amplifying progenitor pool in the oral epithelium ^12^, and, in the case of colon, with the emergence of precancerous stem-cell expansions that precede polyp formation and colorectal cancer ^71^. Of note, in the case of the normal colonic mucosa, EpiDiff displayed higher sensitivity than stemTOC (a mitotic clock), highlighting how it could complement mitotic clocks for cancer risk prediction ^49^. Indeed, it is worth noting that both EpiDiff and mitotic clocks like stemTOC are both strongly enriched for PRC2 targets, yet their CpG overlap is zero with patterns of change in cancer that are opposite: stemTOC displays universal age-acceleration in cancer due to the increased proliferation rate, whilst EpiDiff’s DI is reduced in cancer due to the loss of differentiation. These insights are important given recent work proposing aging and rejuvenation biomarkers that average over PRC2 targets ^72^, thus without considering that there are different functional PRC2 subsets. In the case of EpiDiff CpGs, those gaining DNAm with differentiation were required to be unmethylated in hESCs and gain DNAm gradually from hESC to fetal tissue, as well as between fetal and adult tissue. In contrast, stemTOC’s CpGs were only required to be constitutively unmethylated across many different fetal tissue-types, with the fetal state thus defining the appropriate ground state for the mitotic clock from which mitotic age is then measured.

In the context of aging, we observed how the DI increases with age in many different cell-types, in most cases reflecting an increase in underlying cellular heterogeneity. This makes EpiDiff suitable as a potential aging biomarker to assess age-related shifts in cell-type composition that are detrimental. For instance, given that the fraction of naïve CD4+ T-cells in blood is one of the strongest predictive markers of good health outcomes ^66^, the observed age-associated increase of DI in naïve CD4+ T-cells likely reflects an increased proportion of more differentiated cells that still express the naïve CD4+ T-cell marker but which are functionally compromised. In line with this, the DI of the naïve CD4+ T-cells was strongly anti-correlated with the estimated purity of the naïve CD4+ T-cell samples whilst simultaneously also being strongly correlated with the estimated mature CD4+ T-cell fraction. This highlights how selecting sorted cells with specific markers across lifespan is inappropriate, as the marker does not guarantee purity across age-groups. For other cell-types (e.g hepatocytes, bronchial epithelial cells, astrocytes), the increased DI with age reflects an epigenetic developmental drift towards alternative lineages, specially towards the mesoderm lineage, which would be consistent with the recently observed mesenchymal drift at the transcriptomic level ^73^. However, besides our drift being epigenetic in nature and not transcriptomic, another important difference is that this previous study discovered the transcriptomic mesenchymal drift by analysing age-related gene expression patterns in adult tissues, with the developmental origin of their drift thus being unclear. Here, by starting out from CpGs implicated in early differentiation of the three main germ-layers, we have found that age-associated developmental drift (with mesenchymal drift being the most prominent one) affects CpGs marking differentiation at the earliest stage of development, and hence not just CpGs implicated in adult tissue differentiation and homeostasis. As such, application of EpiDiff to many DNAm datasets profiling sorted cell-types has led to important new insights into the nature of epigenetic aging. Whilst we have no evidence that this DNAm-drift at germ-layer specific CpGs is functionally significant, it is a striking discovery that CpGs implicated in fetal differentiation are more likely to change with age than randomly selected CpGs. Thus, it would be interesting to explore if these developmental DNAm-drifts are eventually linked to downstream functional changes in older cells, providing an epigenetic link to the observed mesenchymal drift at the transcriptional level^73^.

Two important technical features of EpiDiff merit further discussion. First, DI is a measure of global differentiation, in effect quantifying not only the extent of differentiation within one given lineage, but the average extent across all main lineages. We showed this by mixing together sorted mature cell-types from distinct lineages, and noting how these mixtures displayed higher DI-values compared to either one of the original cell-types. This likely explains why differentiated bulk-tissue samples displayed marginally higher DI-values compared to corresponding sorted samples. Similarly, buccal swabs from newborns displayed higher DI values than hematopoietic progenitor cells derived from the bone marrow, likely due to the buccal swabs containing a mix of immature epithelial and immune cell-types. Far from being a limitation, the fact that the DI is sensitive to the underlying cell-type heterogeneity, is a major strength, since it is this feature that allowed the aforementioned discovery of an age-related epigenetic mesenchymal drift. A second technical strength of EpiDiff is that it yields a continuous index on an absolute scale, facilitating direct comparisons between studies and platforms. For instance, we observed how foreskin fibroblast, ovary and testis samples from independent studies were correctly predicted to contain multipotent or progenitor-like cells. Underpinning the ability of EpiDiff to quantify differentiation on an absolute scale, we observed EpiDiff to also display a relatively high resolution. For instance, in a DNAm dataset of bone-marrow derived sorted cells profiling hematopoietic stem cells, multipotent progenitors and downstream lineage-specific progenitors, EpiDiff revealed a small but statistically significant increase of differentiation with the level of commitment. This high resolution was also evident in the ability of EpiDiff to discriminate iPSCs and hESCs according to their capacity to differentiate into the endoderm lineage. The high resolution of EpiDiff also underpins its applicability to sparse single-cell DNAm (snm3C-Seq3) data, correctly predicting an increased differentiation from brain stem-cells (radial glia) along the various neuronal and glia lineages during mid to late gestation, infancy and adulthood. However, we acknowledge that high sparsity of single-nuclei/cell DNAm data could compromise the accuracy of EpiDiff on an absolute scale, since the validations on single-nuclei were done on a relative scale.

We acknowledge other limitations. One potential caveat is that our training set was exclusively made up of Illumina 450k DNAm data. Although this helped avoid the challenge of having to merge DNAm data from different technologies together, it casts a smaller net to identify differentiation-associated CpGs. However, there are a number of other reasons why the training process benefits from considering 450k data as opposed to EPIC or WGBS. First, we required a dataset profiling not only cells from variable differentiation states (pluripotency, multipotency and adult differentiated) but also a dataset that profiled many different types of fetal and matched adult tissues representing all three main germ-layers (endoderm, mesoderm and ectoderm). Most of the current resources providing this data have been generated with the 450k platform. Second, most of the currently available DNAm data has been generated with the 450k technology. Since we want EpiDiff to be as widely applicable as possible, it makes more sense to have trained on 450k CpGs, because training on EPIC arrays would not have rendered the predictor applicable to 450k data: whilst EpiDiff is applicable to EPIC data, owing to the fact that over 90% of the 450k probes are present on EPIC platforms, only about 50% of EPIC probes are present on the 450k platform. Third, the successful validations of EpiDiff in EPIC and single nuclei DNAm data illustrate how starting out from 450k arrays is not a practical limitation. Nevertheless, we acknowledge that improved versions of EpiDiff could be built using future EPIC, WGBS or single-nucleus DNAm data. A second potential caveat is that our training set was composed from data derived from different studies, which could lead to potential batch effects affecting our feature selection strategy. However, we rigorously assessed that batch effects can rarely explain absolute DNAm differences larger than 0.3 or 0.5, which are the thresholds we used for selecting CpGs implicated in fetal and adult differentiation, respectively. Most importantly, our feature selection strategy required that differentiation-CpGs should display significant DNAm changes in both fetal and adult differentiation. Since our starting set of CpGs contained on the order of only 100,000 CpGs and the probability of a batch effect displaying a 0.5 shift or larger was typically only around 1e-5, this means that our selected 1298 diff-CpGs are unlikely to contain false positives. The strong validation performance in the blinded model selection and other independent datasets further demonstrates that EpiDiff is not overfitted to the training data.

In summary, EpiDiff is a unique, robust and accurate tool for predicting the level of differentiation of a DNAm-sample. As demonstrated here, EpiDiff is a versatile tool that can be broadly applied to many different fields, including cellular rejuvenation, regenerative medicine, aging and disease risk prediction. EpiDiff is freely available as an R-package from https://github.com/aet21/EpiDiff.

## Methods

### Construction of EpiDiff and the Differentiation Index (DI)

We built a training DNAm dataset comprising 146 Illumina 450k profiles from various sources including 25 hESC, 37 fetal tissue and 23 adult tissue samples from the Stem-Cell-Matrix Compendium-2 (SCM2) ^40^, 15 cord blood CD34+ samples from Aranyi et al ^74^, 34 fetal tissue samples from Slieker et al ^75^ and 12 adult myeloid samples from Reinius et al ^76^. In total, the DNAm dataset used for feature selection and training consisted of 25 hESC samples representing pluripotency, 86 fetal tissue samples from 14 fetal tissue types representing multipotency, and 35 adult tissue samples from 14 adult tissue-types, representing the fully differentiated state. We also assembled an independent validation (a model-selection set) for optimizing the parameters of the machine learning predictor. This model selection set comprised a total of 180 Illumina 450k profiles, representing pluripotency (29 iPSC samples from SCM2 and 13 hESC samples from Butcher et al ^41^), multipotency (15 CD34+ hematopoietic stem-cell and multipotent progenitor samples from Jung et al ^43^ and 44 buccal swabs from newborn infants ^42^) and fully differentiated states (28 sorted adult differentiated samples from Moss et al ^44^ and 51 adult differentiated bulk tissue samples from Slieker et al ^45^).

Feature selection on the training set was carried out by focusing on 450k CpGs that mapped to within 1500bp upstream of the transcription start site (TSS) or 1^st^ exon of genes, because these are more likely to be of regulatory potential ^77^. This resulted in 100,657 CpGs. We then focused on the subset of 52,433 CpGs that were unmethylated (DNAm beta-value < 0.2) in each of the 25 hESC samples. For each fetal-tissue type, we then performed a differential DNAm analysis using limma to find CpGs that become significantly hypermethylated in that fetal-tissue type compared to the hESC-state. Significance was defined by Benjamini-Hochberg FDR < 0.05 and requiring the average DNAm difference to be larger than 0.3 (justification of this threshold is given in next subsection). A similar differential DNAm analysis was performed for each adult tissue-type comparing to hESCs, but using a larger 0.5 threshold for the average DNAm difference (justification of this threshold is given in next subsection). These DM-analyses resulted in 655 and 638 CpGs displaying significant hypermethylation in at least one fetal-tissue-type and at least one adult-tissue-type respectively, with an overlap of 416 hypermethylated differentiation-CpGs. An analogous analysis was performed for CpGs that were fully methylated (DNAm >0.8) across all 25 hESC samples. In this case, the DM-analyses selected for CpGs that were significantly hypomethylated in at least one fetal and at least one adult-tissue type (using as before FDR<0.05 and 0.3 and 0.5 thresholds on the average DNAm difference for fetal and adult tissue, respectively). In the case of CpGs losing DNAm with differentiation we identified a total of 1877 and 1226 CpGs in the comparisons of hESCs to fetal and adult tissue, respectively, with an overlap of 882 hypomethylated differentiation CpGs. The final set of 416 hypermethylated and 882 hypomethylated differentiation CpGs represent the input to the Random Forest (RF) predictor ^39^. In detail we applied the randomForest function of the *randomForest* R-package to the 1298 differentiation CpG x 146 sample matrix of the training set to learn different RF-predictors parameterized by the number of trees (ranging from 100 to 500, in units of 50, i.e. a total of 9 RF-predictors). Training was performed to predict differentiation state as a categorical variable (pluripotency=0, multipotency=1, differentiated=2). Each of these RF-predictors was then applied to the model selection set of 180 samples. For each sample in the model selection, a given RF-predictor yields probabilities that the given sample is pluripotent (0), multipotent (1) or differentiated (2). Class assignment was done using a maximum probability criterion. However, we also aimed to build a continuous differentiation index (DI). To this end, for each sample *s,* we defined a DI as follows:

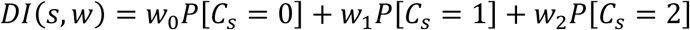

where *w* = (*w*_0_, *w*_1_, *w*_2_) are weight parameters to be optimized and where *P*[*C_s_* = *i*] is the probability from the RF-predictor that sample *s* belongs to differentiation class *i (i=0,1,2).* Of note, 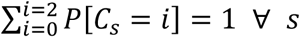. For convenience, we require that the DI index should be 0 when *P*[*C_s_* = 0] = 1 and that the DI index should be maximal (i.e. 1) when *P*[*C_s_* = 2] = 1. Hence, we required that *w*_0_ = 0 and *w*_2_ = 1 are fixed. Besides the number of trees in the RF predictor, the only other free parameter to optimize is *w*_1_, which we allowed to vary between 0.1 and 0.5 in units of 0.025. We did not consider values larger than 0.5 because such high values would lead to worse separability between multipotent and fully differentiated states. Hence, overall, each RF-predictor was parameterized by the number of trees and the *w*_1_ parameter. Performance in the blinded model selection set was carried out by computing the DI-index of each sample and then computing the classification error of each differentiation state. For instance, for the pluripotent state, the error rate was defined as the fraction of true pluripotent samples whose predicted DI values were larger than the minimum DI value across all multipotent samples. We then took the average error-rate over the error-rates of each differentiation state, resulting in an error-matrix defined over the number of trees and *w*_1_ values. The optimal number of trees (=400) and *w*_1_ (=0.25) values were determined by inspection and visualization of this error matrix. This defined the final optimal RF-predictor, which we call EpiDiff.

### Justification of effect size thresholds to avoid confounding by cell-division

When selecting CpGs that are involved in differentiation, we perform a differential DNAm analysis between the pluripotent, multipotent and differentiated states. However, this comparison is in theory confounded by cell-division, since DNAm can change due to errors in DNAm maintenance following cell-division ^48,78^, so that multipotent and differentiated states will have accrued more of these DNAm changes. In order to disentangle DNAm changes associated with cell-division from those associated with differentiation, we impose thresholds on the effect sizes (i.e. difference in DNAm between any two differentiation states), because the changes associated with cell-division are to a large extent stochastic or quasi-stochastic ^25,79^ and hence are much smaller than those typically associated with differentiation ^80^. Indeed, by studying the DNAm effect sizes of CpGs that change with age in a sorted cell-type like monocytes or CD4+ T-cells ^81^, we can obtain highly conservative upper bounds on how much the DNAm level of individual CpGs could change as a result of cell-division. We stress that these are very conservative estimates because age-associated DNAm changes in immune cell-types are not just caused by underlying changes in cell-division, but also result from other factors. Thus, we estimated effect-size distributions of age-DMCs for monocytes and CD4+ T-cells using the corresponding large Illumina 450k DNAm dataset from the MESA-study ^81^, comprising 1199 monocyte and 212 CD4+ T-cell samples from a wide age-range (range in monocytes: 44-83, range in CD4T: 45-79). Specifically, these effect sizes are the change in DNAm over a 1-year period (the slope of a linear regression). We then calculated the change in DNAm over an 80-year period, which is again conservative, as the adult primary samples from the SCM2 training data are younger. Thus, for each cell-type we obtain a distribution of effect sizes (typical change in DNAm over an 80-year period) for all Illumina 450k probes passing QC (482,091 CpGs for monocytes and 482,127 CpGs for CD4T-cells). From this distribution we then estimated the probabilities that a randomly selected CpG would display an effect size larger than a given threshold. At a threshold of 0.5, this probability was as low as 2e-6 or 4e-5 for monocytes and CD4+ T-cells respectively. Thus, given that our feature selection differential DNAm analysis was focused on 100,567 CpGs, this means excellent control of a type-1 error rate. When comparing pluripotent samples to multipotent fetal tissue, we relaxed the threshold to 0.3, for which the corresponding probabilities were 6e-5 and 4e-4, for monocytes and CD4+ T-cells respectively. Because our effect size distribution highly conservative, this choice of threshold also leads to excellent control of the type-1 error rate. Thus, based on these stringent effect size thresholds we can remove any CpGs whose differences in DNAm between differentiation states are the result of increased cell-division. Of note, although the mitotic age effect size distributions were derived from only 2 cell-types (monocytes and CD4+ T-cells), these distributions should be similar for most cell-types that turn over at similar rates and will be overestimates for cell-types that don’t turn over as fast (e.g. lung, neurons).

### Implementation and validation of EpiDiff in independent DNAm datasets

EpiDiff was validated in a large number of independent DNAm datasets, representing variable differentiation states and technologies, including Illumina 450k, Illumina EPICv1, snmCT-Seq and snm3C-Seq3. Irrespective of technology, not all of the 1298 differentiation CpGs making up the EpiDiff predictor may be present in any given dataset. In the case of Illumina DNAm platforms, any missing CpGs were imputed using uninformative mean DNAm values, as computed over the whole data matrix. Thus, the missing CpGs have zero variance and can’t bias patterns in relation to class assignments. In the case of single-cell DNAm data, the 1298 EpiDiff CpGs were mapped to corresponding CpGs in the single-nuclei data. For those EpiDiff CpGs not covered in the single-cell data, we used the spatially most proximal CpG as long as it mapped to within 1kb of the CpG. Of note, because we imputed the low coverage sites in the single-nuclei DNAm data, no further imputation was needed when applying EpiDiff.

### Gene Set Enrichment Analysis (GSEA)

We performed GSEA on the set of 416 hypermethylated and 882 hypomethylated differentiation CpGs which were used as input to the Random Forest (RF) predictor. GSEA was performed in two different ways. One method applied the ebGSEA R-package ^37^ using the Fisher one-tailed enrichment test in conjunction with the Molecular Signatures Database (MSigDB, 2023v2) ^82^. For this analysis, CpGs which already map to regulatory regions (TSS200, TSS1500, 1^st^ Exon), were mapped to NCBI gene identifiers, and the unique genes taken forward for GSEA using ebGSEA. The second method applied the missMethyl R-package ^38^ directly to the CpGs, accounting for differential probe representation across genes.

### Definition of endoderm, mesoderm and ectoderm specific EpiDiff CpGs

From the 1298 EpiDiff CpGs, we identified subsets that were endoderm, mesoderm or ectoderm specific. This was done separately, for the 882 hypomethylated and 416 hypermethylated EpiDiff CpGs. Focusing on the hypomethylated ones, we first assessed in which of the 14 fetal tissue-types the given CpG was significantly hypomethylated (using the same criteria as described in the previous subsection). This defined a binary matrix over the 882 CpGs and 14 fetal tissue-types, with 1 indicating significant hypomethylation, 0 indicating no significant change. Fetal tissue-types were divided into endoderm specific (stomach, liver, lung and pancreas), mesoderm specific (muscle, cord-blood, spleen, heart and kidney) and ectoderm (brain, tongue, adrenal). We note that although tongue and adrenal are of mixed ecto-mesoderm nature, that we placed them in the ectoderm class. For each CpG, we then counted the number of mesoderm, endoderm and ectoderm fetal tissue-types where this CpG was significantly hypomethylated. Subsequently, we defined endoderm-specific hypomethylated CpGs as those significantly hypomethylated in at least one endoderm tissue-type, but not changing in all of the mesoderm and ectoderm tissue-types. We used similar definitions for ectoderm and mesoderm specific hypomethylated CpGs. An exact analogous procedure was carried out to define endoderm, mesoderm and ectoderm-specific hypermethylated EpiDiff-CpGs. All of this resulted in 108 and 139 mesoderm-specific CpGs that lose and gain DNAm with fetal mesoderm differentiation, respectively. Likewise, it resulted in 68 and 71 ectoderm-specific CpGs that lose and gain DNAm with fetal ectoderm differentiation. And finally, 68 and 2 endoderm-specific CpGs that lose and gain DNAm with endoderm fetal differentiation.

### Preferential epigenetic drift at EpiDiff CpGs

For a DNAm dataset of a given sorted cell-type, we assessed if the germ-layer specific EpiDiff CpGs display preferential epigenetic deregulation with age. This was done by first computing age-associated t-statistics and P-values for all CpGs in the dataset, adjusting for potential confounders, which typically included sex, race and technical factors (e.g. Chip-effects). Which confounders to adjust for was decided on a cohort by cohort basis, using a SVD to assess which factors the top principal components (singular vectors) correlate with. Adjustment for multiple-testing was performed using the q-value method ^83^ and statistical significance declared if FDR<0.05. For each dataset, we then calculated a null probability of a CpG significantly changing with age, as well as null probabilities of a CpG significantly hypermethylated (UP) or hypomethylated (DN) with age. For each set of germ-layer specific CpGs we then counted the number of CpGs hypermethylated and hypomethylated with age. Because the germ-layer specific CpGs can become hypermethylated or hypomethylated with differentiation, this resulted in a 2 x 2 matrix counting the number of germ-layer specific CpGs changing with differentiation and age. For each of these numbers we then computed an observed to expected ratio (OER), as well as a P-value from a one-tailed binomial test using the precomputed null probabilities as described above. If the total number of germ-layer CpGs was too small (e.g. for endoderm-hypermethylated CpGs we only had 2), we used a normal approximation.

### Application of EpiDiff to developing human cortical organoids

When applying EpiDiff to the Gordon dataset ^56^, 5.5% of missing CpGs were imputed with the mean value of the DNA methylation matrix. Thus, these CpGs display zero variance and should not bias the estimation of DI. We then computed the Pearson correlation between DI and differentiation day, separately for each of the 4 individuals. To assess if the differentiation state of the organoids at the end of the experiments are comparable to those of adult pre-frontal cortex, we compared the DI of the organoids at the last two timepoints of each donor (n=8 samples) to the DI of 226 adult prefrontal cortex samples, as derived by applying EpiDiff to the Jaffe et al^84^ dataset. This comparison was performed using a one-tailed Wilcoxon rank sum test.

### Application of EpiDiff to human frontal cortex and hippocampus cell lineages across developmental stages

We applied EpiDiff onto the high-resolution developing brain cell lineage dataset from Heffel et al ^57^ to explore if DI tracks cell lineage development. When EpiDiff was applied to the cell type pseudo-bulk samples in this dataset, missing CpGs (20.3%) were imputed with the mean value of the DNA methylation matrix. We then overlayed the estimated DI-values onto the developmental hierarchical tree, with nodes in the hierarchy representing cell types from specific brain regions and developmental stages. For each node in this tree, we assigned the DI value from the corresponding pseudo-bulk sample (if only one exists); for nodes with 2 pseudo-bulk samples (e.g. for glia subtypes like astrocytes derived from both prefrontal cortex and hippocampus), we assigned the mean DI of the corresponding samples. These DI values were then visualized using a gradient color scheme, with cell-type pseudobulk samples (nodes) not passing previous quality control criteria rendered in grey. The Sankey diagram was generated using the ggsankey R package. We also examined whether the DI increases across developmental stage. For excitatory neurons in the prefrontal cortex, glial cells from both the prefrontal cortex and hippocampus, CGE-derived inhibitory neurons, and MGE-derived inhibitory neurons, P-values were derived from linear regression of DI against developmental stage. For excitatory neurons in the hippocampus the pattern was clearly non-linear, hence in this case, the P-value was obtained from a one-tailed Wilcoxon test comparing the DI values of the earliest stage (2T) to all other later stages.

### *In-silico* mixture analysis of sorted and single cells

For DNAm datasets profiling sorted cell-types we only used donors contributing a sample for each cell-type, in order to then generate mixtures by mixing together the cell-types from the same donor. This is important to remove genetics as a confounder. Thus, assuming that there are *n* donors each contributing *m* cell-types, we generated for each pair of cell-types *t_1_* and *t_2_*, *n* in-silico mixtures:

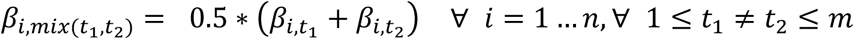

In the case of sorted immune cell-types we used the data from Bell et al (*m=*6 mature immune cell-types, so 15 pairwise mixtures with *n*=114) and BLUEPRINT (*m*=3 immune cell-types, 2 mature ones, monocytes and neutrophils, and naïve CD4+ T-cells, with *n*=139) ^60,61^. In the case of brain, we used the sorted data from Walker et al ^62^ comprising *n*=114 donors each contributing a neuron, oligodendrocyte and astrocyte sample (*m*=3). For each pair of cell-types, we then compared the DI-values of the mixed samples to the DI’s of the original cell-types, using a paired Wilcoxon rank sum-test. In the case of the 3 brain cell-types (Walker et al), we also generated mixtures of the 3-cell-types, comparing the DI of the 3 cell-type mixtures to those of 2 cell-type mixtures.

For Chien et al’s cell-type-level summarized snmCT-seq dataset, comprising deep layer excitatory neurons, intermediate layer excitatory neurons, upper layer excitatory neurons, CGE-derived inhibitory neurons, MGE-derived inhibitory neurons, astrocytes, microglia, and oligodendrocytes, mixtures ranging from 2 to all 8 cell types were generated. We employed linear regression to evaluate the relationship between DI and the number of constituent cell types in the mixture.

Of note, the fraction of missing EpiDiff CpG sites in these datasets were: BLUEPRINT, 5.3%; Bell, 3.2%; Walker, 8.9%; snmCT-seq dataset, 0.2%. In the latter case, the fraction is lower because when EpiDiff CpGs did not match by exact genomic position, we used a 1000bp window to select the closest proxy CpG. In all cases, missing/unassigned CpG sites were imputed with the mean value of the DNA methylation matrix to avoid biasing the DI-estimation.

### Meta-Analysis of DI in glial cells and age

Here we used the cell-type level summarized snmCT-Seq dataset from Chien et al ^63^, a study of aging in neurons and glia. EpiDiff was applied on the glial subtypes. Missing EpiDiff CpG sites (0.2%) were imputed with the mean value of the DNA methylation matrix to avoid biasing the DI-estimation. To enhance statistical power, meta-analysis was employed to integrate findings across the glia subtypes (astrocytes, microglia, and oligodendrocytes). In more detail, for each glial subtype, we first fitted multivariate linear regression models with DI as the dependent variable and age group (old vs. young) as the independent variable, adjusting for sex. Effect sizes, standard errors, t-statistics, and P-values were extracted from each model. We then performed meta-analyses using inverse variance weighted methods under both fixed-effect and random-effects assumptions, implemented with the metagen function in the meta R-package^85^.

### Processing and normalization of Illumina DNAm data

Unless specified otherwise, Illumina DNAm datasets were processed and normalized as follows. For datasets with available idat files, the preprocessRaw and ratioConvert functions from the minfi package (version 1.56.0) were employed to convert Red and Green channel intensities into a matrix of Beta values ^86^; for datasets where raw idat files were unavailable, processed beta-valued matrices were utilized. The getAnnotation function of the minfi package was used to identify and exclude probes located on sex chromosomes, and the dropLociWithSnps function was employed to remove probes with potential SNPs at the CpG site or single base extension site. Additionally, probes identified as having cross-reactivity were excluded. Beta values with poor detection quality (P-values > 0.01) were masked as missing. The handling of missing values depended on sample size: for datasets with >100 samples, probes with a coverage fraction less than 0.9 were discarded and remaining missing values imputed using the k-nearest neighbors method (impute.knn, k = 5, impute package v1.84.0) ^87^; conversely, for datasets with <100 samples, probes containing any missing values were excluded. Finally, to correct for probe design bias (Type II vs. Type I), the BMIQ algorithm was applied to the processed data ^88^.

### Training set for building EpiDiff

We assembled a training set of 146 samples all generated with Illumina 450k platform. These 146 samples comprised 25 hESCs from the SCM2 compendium ^40^, 86 multipotent (fetal) samples (including 37 fetal tissue samples from SCM2, 15 cord-blood CD34+ samples from Aranyi et al ^74^ and 34 fetal tissue samples from Slieker et al ^75^) and 35 adult differentiated samples (including 23 from the SCM2 and 12 sorted myeloid adult samples from Reinius et al ^76^. The SCM2 dataset in processed form was downloaded from GEO GSE31848. Probes with less than 5% missing values were kept and imputed using impute.knn (k=5). Subsequently we applied BMIQ to adjust for type-2 probe bias. In the case of cord-blood CD34+ samples, raw idat files were downloaded from GEO GSE72867 and processed with minfi and subsequently normalized with BMIQ. Processed normalized data from Slieker et al was downloaded from GEO GSE56515 and further normalized with BMIQ. The 450k data from Reinius et al consists of sorted immune cell samples, was downloaded from GEO GSE35069 (raw idat files), and subsequently processed with minfi and normalized with BMIQ. Feature selection and training was restricted to 100,657 common probes that map to either 1^st^ exon or upstream of the TSS up to a distance of 1500bp. The 37 fetal-tissue samples from SCM2 comprised 3 adrenal, 4 brain, 4 heart, 5 kidney, 4 liver, 5 lung, 3 spleen, 5 stomach, 2 thymus and 2 tongue samples. The 23 adult-tissue samples from SCM2 comprised 2 adipose, 2 adrenal, 2 bladder, 2 blood, 1 brain, 1 heart, 2 lung, 2 lymph-node, 2 pancreas, 2 skeletal muscle, 2 spleen, 1 stomach and 2 ureter samples. The 34 fetal-tissue samples from Slieker et al comprised 9 adrenal, 8 amnion, 9 muscle and 8 pancreas samples.

### Model selection (validation) set for optimizing parameters

To select optimal parameters, we optimized performance in a model selection set of 180 samples comprising 5 independent DNAm datasets and representing 42 pluripotent samples (29 iPSC samples from the SCM2 compendium ^40^ and 13 hESC samples from Butcher et al ^41^), 59 multipotent samples (44 buccal swabs from newborns ^42^ and 15 CD34+ hematopoietic multipotent cells from Jung et al ^43^) and 79 adult tissue samples (28 sorted samples from 11 different tissue-types from the DNAm-atlas of Moss et al ^44^ and 51 samples from 9 tissue-types from Slieker et al ^45^). The 13 hESC samples from Butcher et al and the 15 CD34+ samples from Jung et al are part of larger DNAm datasets, whose processing and normalization procedure is described further below. The buccal swabs from 44 newborns is an EPICv1 DNAm dataset that derives from Kocher et al ^42^. Raw idat files were downloaded from GEO GSE229463, processed and normalized as described earlier. The DNAm-atlas from Moss et al contains 450k, EPICv1 and EPICv2 profiles and raw idat files were downloaded from GEO GSE122126, and subsequently processed with minfi and normalized with BMIQ. We processed each platform separately, finally merging the 3 datasets together for a total of 449,156 probes and 28 sorted samples (10 pancreatic, 3 adipocyte, 3 lung epithelial, 3 colon epithelial, 3 hepatocyte, 3 cortical neurons, 2 vascular endothelial and 1 leukocyte). The adult tissue samples from Slieker et al is an Illumina 450k dataset for which the processed normalized data was downloaded from GEO GSE48472. Data was subsequently adjusted for type-2 probe bias using BMIQ. Of the original 56 samples, we removed 5 hair-follicle samples, due to these samples containing multipotent cells, leaving a total of 51 samples (11 blood, 5 buccal,5 liver, 6 muscle, 6 omentum, 4 pancreas, 5 saliva, 6 subcutaneous fat and 3 spleen).

### DNAm profiling of cell-lines undergoing population doublings

The Illumina EPICv1 cell line data ^46^ was downloaded from GEO under accession number GSE197512. Raw idat files were processed with *minfi* and BMIQ normalized, resulting in a normalized DNAm data matrix defined over 843,298 probes and 182 “baseline-profiling” samples. A total of 7 human cell lines were used, including AG06561 (skin fibroblast), AG11182 (veil endothelial), AG11546 (iliac vein smooth muscle), AG16146 (skin fibroblast), AG21839 (neonatal foreskin fibroblast), AG21859 (foreskin fibroblast) and AG21837 (skin keratinocyte). Population doublings ranged from 7 to 92. The number of samplings per cell-line ranged from 16 (AG16146) to 40 (AG21859).

### DNA methylation datasets of sorted immune cell-types

#### MESA study (GSE56581)

The Illumina 450k set from Reynolds et al (MESA study) ^81^ contributed 1199 monocyte and 212 CD4+ T-cell samples, after QC. We used the normalized DNAm data as described by us previously ^80^.

#### BLUEPRINT (BP) ^61^

We obtained 139 monocyte, 139 naïve CD4+ T-cell and 139 neutrophil samples from 139 individuals. We used the normalized DNAm data as described by us previously ^80^.

#### Bell et al (GSE224807)

The Illumina HM450k dataset from Bell et al. ^89^ profiling normal sorted blood cells was obtained from the GEO database under the accession number GSE224807. This cohort comprises 789 samples. Chronological age information is not available for this dataset. The samples encompass 123 CD4+ T cells, 124 CD8+ T cells, 129 B cells, 129 monocytes, 122 natural killer (NK) cells, 126 granulocytes with 114 donors each contributing 6 cell-type samples.

#### Salas et al. (GSE167998) 51

This dataset comprises EPICv1 profiles of 12 sorted immune-cell subsets, totaling 56 sorted samples: 6 basophils (Baso), 6 memory B-cells (Bmem), 4 naive B-cells (Bnv), 4 memory CD4+ T-cells (CD4Tmem), 5 naive CD4+ T-cells (CD4Tnv), 4 memory CD8+ T-cells (CD8Tmem), 5 naive CD8+ T-cells (CD8Tnv), 4 eosinophils (Eos), 5 monocytes (Mono), 6 neutrophils (Neu), 4 natural killer (NK) cells, and 3 regulatory T-cells (Treg). This dataset had previously been processed and normalized by us in Luo et al. ^66^.

#### Sorted cord-blood cell-types

We used a merged normalized Illumina 450k DNAm dataset of 7 cord blood cell-types (11 nucleated Red Blood Cell samples, 28 B-cell samples, 29 CD4+ and 20 CD8+ T-cell samples, 22 NK-cell, 24 granulocyte and 32 monocyte samples), which underwent quality control and normalization as described by us previously ^50^.

### DNA methylation datasets of other sorted cell-types

#### Brain (Neurons, astrocytes, oligodendrocytes and microglia)

The EPICv1 dataset from Walker et al. ^62^ profiling sorted prefrontal cortex cells from normal individuals was obtained from the GEO database under the accession number GSE279509. This cohort comprises 751 samples, with ages ranging from 18 to 108 years. Specifically, the sorted cells include 164 Double-(astrocytes), 12 IRF8+ (microglia), 182 NeuN+ (neurons) and 168 Sox10+ (oligodendrocytes), with a total of 114 donors contributing a sample for each of these 4 cell-types.

#### Liver Hepatocytes

We analyzed an Illumina EPICv1 DNAm dataset comprising 56 hepatocyte cultures. Raw idat files were downloaded from GEO under accession number GSE123995, and processed using minfi. We only selected probes with detected values (P<0.01) across all samples, resulting in a dataset of 856,856 CpGs and 56 samples. One sample was removed due to displaying an abnormal DNAm density profile. The remaining dataset of 55 samples was then normalized using BMIQ ^90^.

#### Bronchial Epithelial Cells (BECs)

This is a joint 450k and EPIC DNAm dataset profiling bronchial epithelial cells from asthmatic cases and controls. Raw idat files were downloaded from GEO GSE201872 and processed using minfi. For both 450k and EPIC DNAm datasets we restricted to probes with detected P-values (P<0.01) across all samples, subsequently merging the two data matrices, resulting in a data matrix of 449,679 probes and 142 samples. This data matrix was then normalized with BMIQ. For the age-analysis, one sample that was unique to one beadchip was removed, as its inclusion would not allow proper adjustment for beadchip effects when correlating DI to age, resulting in 141 samples (46 controls and 95 asthmatic cases).

#### Pulmonary Endothelial Cells (PECs)

This is an Illumina 450k DNAm dataset of 18 pulmonary endothelial cell samples. Raw idat files were downloaded from GEO GSE84395 and processed with minfi. Only probes with detection P-values < 0.01 across all 18 samples were used, resulting in a data matrix of 484,081 CpGs and 18 samples, which was subsequently normalized with BMIQ.

#### Bone Marrow Hematopoietic Cell-types

This is an Illumina 450k DNAm dataset of 74 samples from Jung et al ^43^. Raw idat files were downloaded from GEO GSE63409 and processed with minfi. Only probes with detection P-values < 0.01 across all samples were used, resulting in a data matrix of 460,726 CpGs, which was subsequently normalized with BMIQ. We restricted analysis to a subset of 30 samples comprising 5 hematopoietic stem-cell (HSC), 10 multipotent progenitor (MPP), 5 megakaryocyte-erythrocyte progenitor (MEP), 5 granulocyte-myeloid progenitor (GMP) and 5 common myeloid progenitor (CMP) samples.

### In-vitro differentiation DNAm datasets

#### Developing human cortical organoid (Gordon et al)

The Illumina EPIC DNAm dataset from Gordon et al^56^ profiled DNAm in a differentiation time course of human iPSCs into human cortical organoids. Data was obtained from the NCBI GEO website under the accession number GSE150122 (https://www.ncbi.nlm.nih.gov/geo/query/acc.cgi?acc=GSE150122). The file “GSE150122_RAW.tar” which contains the IDAT files was downloaded and processed with minfi package. Probes with NAs (defined by P > 0.01) were discarded. The filtered data was subsequently normalized with BMIQ, resulting in a normalized data matrix for 856809, probes and 52 samples. Samples were collected at different timepoints (days of differentiation) from 5 donors, but only 4 donors had samples collected at more than one timepoint, resulting in 51 samples.

#### Differentiation of Mesenchymal Stem Cells into Hepatocytes

This is an Illumina EPICv1 DNAm dataset from Takashina T et al ^55^ consisting of 4 samples. We downloaded the processed data from GEO GSE205546. We selected probes with detected values (P< 0.01) across all samples. The resulting data matrix of 859,841 probes and 4 samples was then normalized with BMIQ. These 4 samples consist of (i) umbilical cord-blood derived mesenchymal stem cells (MSCs), (ii) MSCs that were treated with NE for 5 days (negative controls) and (iii+iv) MSCs treated with HNF1A, HNF3G, HNF4A and GATA4 (4F) for 3 and 5 days, respectively.

### Single-nuclei DNAm data

#### Human frontal cortex snmCT-seq dataset

The snmCT-seq dataset from Chien et al^91^, profiling DNAm in human frontal cortex cells, was obtained from the NCBI GEO website under the accession number GSE247988 (https://www.ncbi.nlm.nih.gov/geo/query/acc.cgi?acc=GSE247988). A total of 20 samples were collected from 11 donors (5 young, aged 23-30 years; 6 old, aged 70-74 years), with 1-2 tissue chunks processed independently per donor. Each sample yielded approximately 3,000 cells. Single-nucleus methylome sequencing files were downloaded for each sample, and cells failing quality control according to the metadata from Chien et al were excluded. Given the limited coverage per single cell, we conservatively retained all sequenced CpG sites for downstream analysis. Mapping of these CpG sites to Illumina Infinium HumanMethylation450K array probes was performed using a two-step strategy: first, probes were matched based on exact genomic coordinates; for probes that could not be uniquely assigned via exact mapping, the nearest CpG site within a ±1,000 bp window was used as a proxy. To mitigate technical noise and obtain a more stable representation of each cell type, we aggregated data by calculating mean DNA methylation values at the cell-type level (8 cell types: deep layer excitatory neurons, intermediate layer excitatory neurons, upper layer excitatory neurons, CGE derived inhibitory neurons, MGE derived inhibitory neurons, astrocytes, microglia cells, oligodendrocytes). After applying coverage thresholds of >0.6 at the sample level and >0.9 at the probe level, the final summarized cell-type dataset comprised 148 samples and 483,277 CpGs. The remaining NAs were imputed with impute.knn (k = 5) ^92^.

#### Human frontal cortex and hippocampus snm3C-seq3 dataset

The snm3C-seq3 dataset from Heffel et al^57^ profiled DNAm in single nuclei from the human frontal cortex and hippocampus across 4 developmental stages: mid gestation (2T:GW18-GW23), late gestation (3T:GW35-GW39), infants (4-7 months) and adults (21-37 year olds). Data was obtained from the NCBI GEO website under the accession number GSE213950 (https://www.ncbi.nlm.nih.gov/geo/query/acc.cgi?acc=GSE213950). A total of 188 cell-type specific pseudobulk data files were downloaded. For each cell type’s pseudobulk data file, the CpGs were filtered by demanding at least 5 read coverage in each sample and by subsequent mapping to Illumina 450K array probes. Applying a threshold of 0.5 for the sample coverage and 0.9 for CpG coverage, this resulted in a data matrix comprising 366,374 CpGs and 130 cell-type pseudo-bulk samples. The remaining NAs were imputed with impute.knn (k=5)^92^. Meta-information about each cell-type was downloaded from https://brain-epigenome.cells.ucsc.edu/.

### Full reprogramming DNAm datasets

#### Butcher

The 450K dataset from Butcher et al ^41^ profiled DNAm in human dermal fibroblasts, endothelial precursors, foreskin fibroblasts and their induced iPSCs. DNAm data was obtained from the NCBI GEO website under the accession number GSE59091 (https://www.ncbi.nlm.nih.gov/geo/query/acc.cgi?acc=GSE59091). The file “GSE59091_unmethylated_methylated.txt.gz” was downloaded and processed. Probes with NAs (defined by P > 0.01) were discarded. The filtered data was subsequently normalized with BMIQ, resulting in a normalized data matrix for 475559 probes and 166 samples.

#### Wang

The 450K dataset from Wang et al ^93^ profiled DNAm in human skin fibroblast, vein endothelial cells and their induced iPSCs. Data was obtained from the NCBI GEO website under the accession number GSE68134 (https://www.ncbi.nlm.nih.gov/geo/query/acc.cgi?acc=GSE68134). The file “GSE68134_non-normalized.txt.gz” was downloaded and processed. Probes with NAs (defined by P > 0.01) were discarded. The filtered data was subsequently normalized with BMIQ, resulting in a normalized data matrix for 461990 probes and 41 samples.

#### Fernandez

The 450K dataset from Fernandez et al ^94^ profiled DNAm in human fibroblasts, LCL and their induced iPSCs was obtained from the NCBI GEO website under the accession number GSE51921 (https://www.ncbi.nlm.nih.gov/geo/query/acc.cgi?acc=GSE51921). The file “GSE51921_signal_intensities.txt.gz” was downloaded and processed. Probes with NAs (defined by P > 0.01) were discarded. The filtered data was subsequently normalized with BMIQ, resulting in a normalized data matrix for 461921 probes and 14 samples.

#### Burrows

The 450K dataset from Burrows et al ^95^ profiled DNAm in human fibroblast and the induced iPSCs was obtained from the NCBI GEO website under the accession number GSE65078 (https://www.ncbi.nlm.nih.gov/geo/query/acc.cgi?acc=GSE65078). The file “GSE65078_RAW.tar” which contains the IDAT files was downloaded and processed with minfi package. Probes with NAs (defined by P > 0.01) were discarded. The filtered data was subsequently normalized with BMIQ, resulting in a normalized data matrix for 484335 probes and 24 samples.

### Partial reprogramming and cellular rejuvenation DNAm datasets

#### Sarkar

The EPIC dataset from Sarkar et al ^58^ profiled DNAm in human fibroblast and endothelial cells and their partially reprogrammed ‘rejuvenated’ cells. For fibroblasts and endothelial cells a total of 4 biological replicates each were performed, resulting in a total of 16 samples: 4 somatic fibroblast and 4 somatic endothelial samples, and 8 rejuvenated samples. Data was obtained from the NCBI GEO website under the accession number GSE142439 (https://www.ncbi.nlm.nih.gov/geo/query/acc.cgi?acc=GSE142439). The file “GSE142439_RAW.tar” which contains the IDAT files was downloaded and processed with minfi package. Probes with NAs (defined by P > 0.01) were discarded. The filtered data was subsequently normalized with BMIQ, resulting in a normalized data matrix for 857,471 probes and 16 samples.

#### Gill

The EPIC dataset from Gill et al ^29^ profiled DNAm in human dermal fibroblast and the partial reprogrammed cells was obtained from the NCBI GEO website under the accession number GSE165180 (https://www.ncbi.nlm.nih.gov/geo/query/acc.cgi?acc=GSE165180). The file “GSE165180_RAW.tar” which contains the IDAT files was downloaded and processed with minfi package. Probes with NAs (defined by P > 0.01) were discarded. The filtered data was subsequently normalized with BMIQ, resulting in a normalized data matrix for 847791 probes and 49 samples.

### Differentiation Capacity (DC) of iPSCs

The Illumina 450k DNAm dataset of Butcher et al ^41^ (GEO GSE59091) was processed as described in the section on full reprogramming DNAm datasets. Here we focused on 116 iPSCs derived from dermal fibroblasts (D-iPSC), 10 iPSCs generated from endothelial precursors (E-iPSC) and 13 hESCs, all of which were characterized in terms of their ability to differentiate into the endoderm lineage. In total, among the 116 D-iPSCs there were 49 and 67 displaying high and low DC, respectively. Among the 10 E-iPSCs, 4 and 6 were categorized as high and low DC, respectively. Of the 13 hESCs, 5 and 8 were of high and low DC, respectively.

### Bulk-tissue DNAm datasets

#### Cord-Blood

We used an Illumina 450k set comprising 98 cord-blood samples from Paul et al ^52^. We used the normalized DNAm dataset as described in Paul et al.

#### Enhanced GTEX

The enhanced GTEX resource is a large multi-tissue EPICv1 DNAm dataset comprising 987 adult samples from 9 tissue-types (colon, lung, testis, prostate, ovary, skeletal muscle, kidney, breast and blood) first presented in Oliva et al ^53^. We used the normalized DNAm data as described in our previous publication ^49^.

#### Skin Dermis and Epidermis

This Illumina 450k DNAm dataset of 78 skin samples (dermis & epidermis) is from Vandiver et al ^96^. We have previously processed and analyzed this DNAm dataset in ^80^.

#### Buccal Swabs

The Illumina 450k DNAm dataset of 790 buccal swab samples from women all aged 53 derives from the NSHD MRC1946 birth cohort. We used the normalized DNAm data, as described in our previous publications ^12,97^.

#### Liver

This is an EPICv1 DNAm dataset of liver samples from Johnson et al ^98^. We downloaded the processed DNAm beta-values and P-values from GEO GSE180474. We then restricted to probes with detection P-value (P<0.01) across all samples, subsequently normalizing the dataset with BMIQ. In this work, we focused only on 210 samples of normal histology. We previously analyzed this DNAm dataset in Tong et al ^99^.

#### Vaginal swabs

This is an Illumina 450k DNAm dataset previously generated and analyzed by us ^100^. We used the BMIQ normalized data as used in this previous study. Here we focused on 73 vaginal swab samples from healthy women.

#### Esophagus

This is an Illumina 450k DNAm dataset from Luebeck et al ^23^ profiling 52 samples from the normal healthy esophagus. We downloaded the processed DNAm data from GEO GSE104707. Subsequently, the data matrix was adjusted for type-2 probe bias using BMIQ. We have previously analyzed this DNAm dataset in ^80^.

#### Stomach

This is an Illumina 450k DNAm dataset from Huang et al ^101^ profiling 191 gastric samples, including 61 normals and 130 intestinal metaplasias. We downloaded the raw idat files from GEO GSE103186 and process them with minfi. We selected probes with coverage of at least 0.99, imputing the rest with the *impute.knn* function from impute R-package. Subsequently, data was normalized with BMIQ, resulting in a DNAm data matrix of 482,975 probes and 191 samples. We have previously analyzed this DNAm dataset in ^80^.

#### Adult prefrontal cortex (Jaffe et al)

We used the 226 prefrontal cortex samples from healthy adults derived from the Illumina 450k dataset from Jaffe et al.^84^. This dataset is available from GEO under accession number GSE74193. DNAm data was normalized and processed as previously described^99^.

#### TCGA datasets

The SeSAMe ^102^ processed Illumina 450k beta value matrices were downloaded from Genomic Data Commons Data Portal (https://portal.gdc.cancer.gov/) with TCGAbiolinks ^103^. We here analyzed 16 cancer types (BLCA, BRCA, COAD, ESCA, HNSC, KIRC, KIRP, LIHC, LUAD, LUSC, PAAD, PRAD, READ, SKCM, STAD, THCA). We did further processing of the downloaded matrices as follows: For each cancer type, probes with missing values in more than 30% samples were removed. The missing values were then imputed with impute.knn (k=5) ^92^. Then, the type-2 probe bias was adjusted with BMIQ ^90^. Technical replicates were removed by retaining the sample with highest CpG coverage. Clinical information of TCGA samples was downloaded from Liu et al ^104^.

#### Erlangen breast cancer

This Illumina DNAm dataset comprises 50 normal breast samples, 42 normal-adjacent and 305 breast tumors, and has been extensively analyzed by us previously ^68^. *Colon mucosa and crypts:* This joint Illumina 450k and EPIC DNAm dataset profiled normal samples from the gastrointestinal system including colon, duodenum, ileum, jejunum and small intestine (Lewis et al.) ^105^. Raw idat files were obtained from GEO under accession number GSE141254 and was normalized as described earlier. Here we focused on mucosa and crypt samples from the colon: 10 crypt and 9 mucosa samples for the 450k experiment, 8 crypt and 3 mucosa samples for the EPICv1 experiment.

#### Colon Cancer Risk

This joint 450k and EPICv1 DNAm dataset profiled normal colon mucosa samples from individuals categorized as of low colorectal risk (no concurrent polyps or cancer), medium risk (concurrent polyps but no cancer) and high risk (concurrent colorectal cancer) ^70^. Raw idat files were downloaded from GEO GSE132804, processed and normalized as described earlier. In total there were 334 samples (128 and 206 profiled with 450k and EPIC platforms, respectively). Of the 334 samples, 105 were of low cancer risk, 128 of medium risk and 101 of high-risk.

## Data availability

The following DNA methylation datasets analyzed in this study are all publicly available from Gene Expression Omnibus (GEO, www.ncbi.nlm.nih.gov/geo/) under the following accession numbers: GSE31848 (SCM2), GSE72867 (Aranyi), GSE56515 (Slieker, fetal tissues), GSE143209 (Reinius), GSE59091 (Butcher), GSE63409 (Jung), GSE229463 (Kocher), GSE122126 (Moss), GSE48472 (Slieker, adult tissues), GSE56581 (MESA study), GSE150122 (Gordon), GSE74193 (Jaffe), GSE213950 (Heffel), GSE224807 (Bell), GSE279509 (Walker), GSE247988 (Chien), GSE197512 (cell-lines undergoing population doublings), GSE167998 (Salas), GSE123995 (Liver Hepatocytes), GSE201872 (Bronchial Epithelial Cells), GSE84395 (Pulmonary Endothelial Cells), GSE205546 (Takashina), GSE68134 (Wang), GSE51921 (Fernandez), GSE65078 (Burrows), GSE142439 (Sarkar), GSE165180 (Gill), GSE213478 (Enhanced GTEX), GSE52980 (Vandiver), GSE180474 (Johnson), GSE104707 (Luebeck), GSE103186 (Huang), GSE69914 (Erlangen breast cancer), GSE141254 (Colon mucosa and crypts), GSE132804 (Colon Cancer Risk). The BLUEPRINT DNAm data of sorted monocytes, neutrophils and CD4+ T-cells is available from EGA under accession number EGAS00001001456. The DNAm dataset of buccal swabs from NSHD MRC1946 birth cohort is available by submitting data requests to; see full policy at http://www.nshd.mrc.ac.uk/data.aspx. TCGA Illumina 450k datasets for BLCA, BRCA, COAD, ESCA, HNSC, KIRC, KIRP, LIHC, LUAD, LUSC, PAAD, PRAD, READ, SKCM, STAD, and THCA datasets are available from the GDC data portal https://portal.gdc.cancer.gov/. The merged dataset of sorted cord-blood cell-types data and the Vaginal swabs data are available upon request to the corresponding author.

## Code availability

EpiDiff is freely available as an R-package from https://github.com/aet21/EpiDiff. This R-package contains a brief user-friendly tutorial.

## Acknowledgements

This work was supported by NSFC (National Science Foundation of China) grants, grant numbers 32370699 and 32570775, and an RFIS grant W2431024. We also thank Stephan Beck and Nir Eynon for comments provided on previous versions.

## Authors contributions

AET conceived the study. AET and HT performed statistical analyses with contributions from XG. AET wrote the manuscript with contributions from HT and XG.

## Competing interests

The authors declare that they have no competing interests.

## Notes

### Competing Interest Statement

The authors have declared no competing interest.

