## Supplementary Information for "DNA methylation-based prediction of differentiation state at cell-type resolution reveals an age-associated epigenetic mesenchymal drift"

### Supplementary Figures

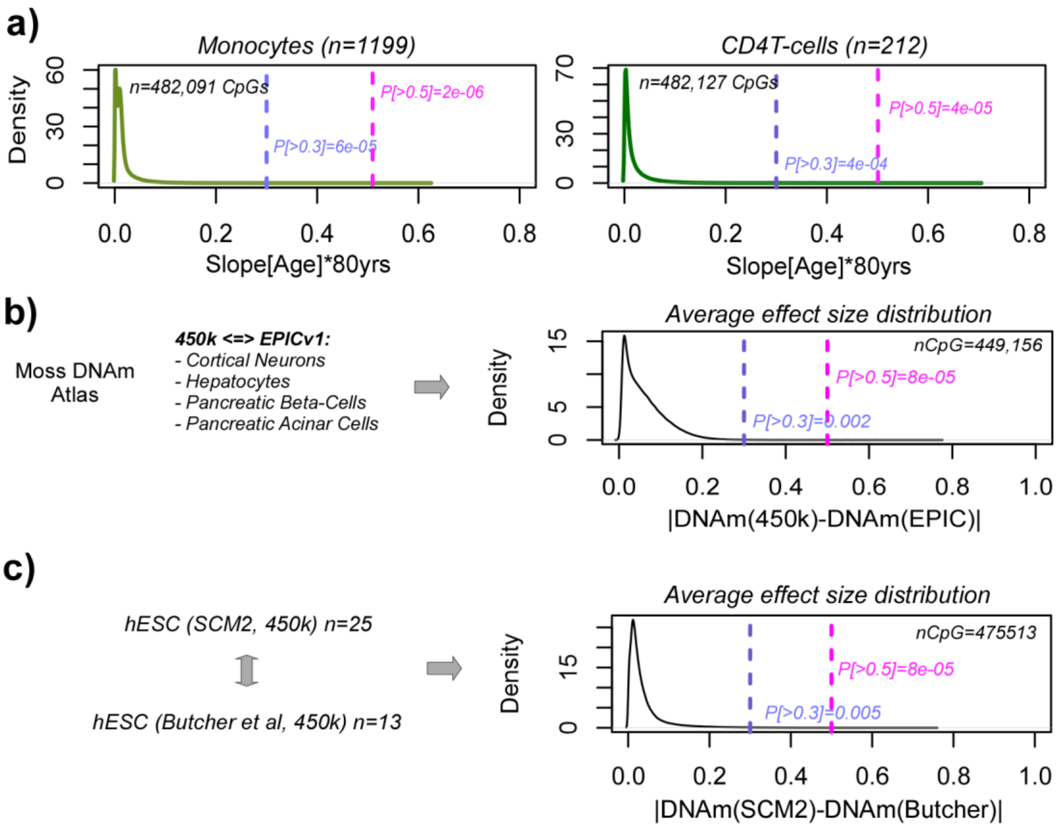


**SI fig.S1: Justification of effect size thresholds. a)** Density plots of age-associated effect sizes of CpGs in two large sorted Illumina 450k datasets from the MESA-study. The effect size is shown as the slope (estimated DNAm change per year) multiplied by 80 years, to get the lifetime effect size. Since the adult tissues from the SCM2 cohort are younger and because mitotic age is only one component of age-associated DNAm changes, this gives a very conservative (stringent) effect size distribution for how big cell-division associated DNAm changes can be. Vertical dashed lines give the probabilities of a CpG displaying effect sizes larger than 0.3 or 0.5. **b)** Using the Moss DNAm-atlas where 4 cell-types were profiled separately with 450k and EPICv1 platforms, we can estimate the effect size distribution of typical batch effects, by computing the absolute DNAm difference between 450k and EPIC samples for each of the 4 cell-types and subsequently averaging over the 4 cell-types, as shown in density plot. Vertical dashed lines give the probabilities of a CpG displaying effect sizes larger than 0.3 or 0.5. **c)** Density plot of absolute differences in DNAm between 25 hESCs profiled in SCM2 and 13 hESCs profiled in Butcher et al, both profiled with HM450k arrays. Vertical dashed lines give the probabilities of a CpG displaying effect sizes larger than 0.3 or 0.5.

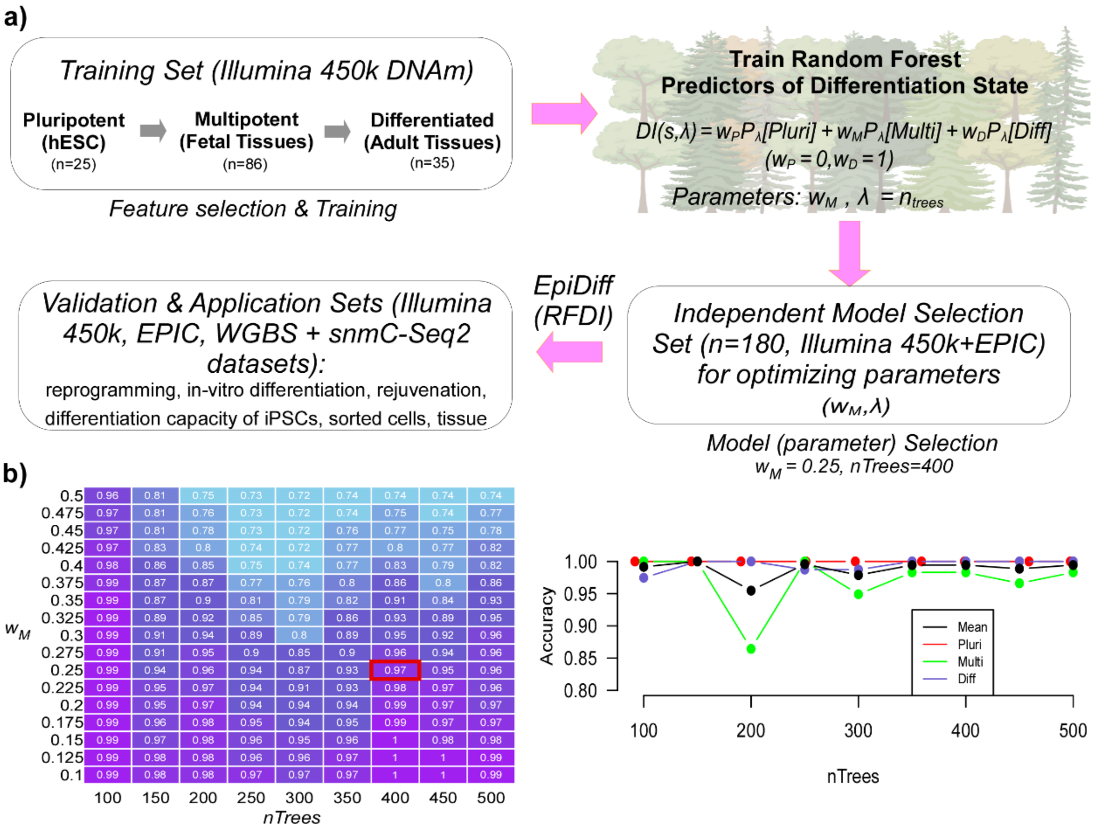


**SI fig.S2: Overall strategy to build and validate EpiDiff. a)** A collection of Illumina 450k DNAm datasets encompassing a total of 146 samples comprising pluripotent, multipotent (fetal) and adult tissue differentiated states was used for feature selection and training a RandomForest (RF) Classifier. This RF-classifier was run in categorical mode with nodesize=1, and returns probabilities of a sample belonging to pluripotent, multipotent and differentiated categories. These probabilities were then combined to build a Differentiation Index (DI) as shown, with one free parameter (weight) $w_M$ . The other free parameter to be optimized is the number of trees in the RF classifier. We then used a collection of independent DNAm datasets encompassing pluripotency, multipotency and adult differentiated states for model/parameter optimization and selection. Finally, this optimized RFDI predictor (called EpiDiff) was validated in independent DNAm datasets. **b)** Optimization landscape heatmap displaying the overall classification accuracies as a function of the two free parameters. To ensure robustness in the context of omic datasets, we favoured solutions with more trees, with nTrees=400 being optimal. We then chose $w_M$*=0.25* over lower $w_M$ values as difference in performance was similar and within statistical error intervals, so as to give more weight to intermediate multipotent state. Scatterplot to the right shows how the accuracy varies per category + overall accuracy for $w_M$*=0.25* as a function of *nTrees.*

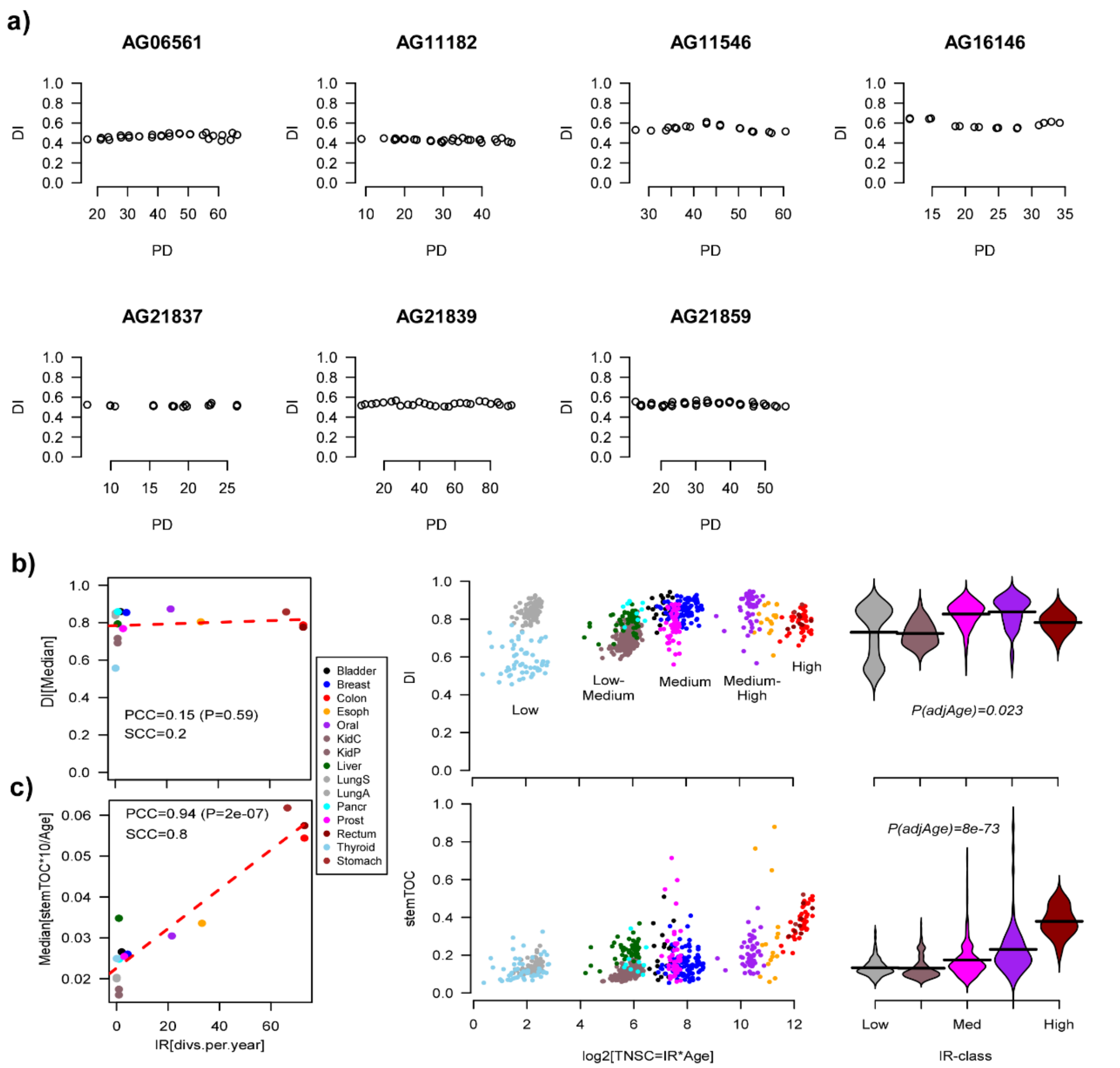


**SI fig.S3: DI is not altered with cell-division. a)** For each of 7 cell-lines (EPIC DNAm data) that underwent populations doublings (PDs), we plot the predicted DI-values against PD. **b)** Left: For all the normal-adjacent tissue samples from the TCGA, we plot the median DI per tissue-type (y-axis) against the intrinsic rate (IR) of stem-cell division of that tissue-type. Pearson and Spearman correlation coefficients (PCC, SCC) are given. For the PCC we also provide the P-value from a correlation test. Red dashed line is the best regression fit. Middle: As left, but now displaying all the samples from each tissue-type, so y-axis now labels the DI and x-axis labels the total number of stem-cell divisions (TNSC) for each sample in a log2-basis. Right: Violin representation of middle panel, with tissue-types stratified into 5 categories: low IR, low-medium IR, medium IR, medium-high IR and high IR. P-value of a linear regression of DI against log2(TNSC) adjusted for chronological age is given. **c)** As a) but for the mitotic clock stemTOC.

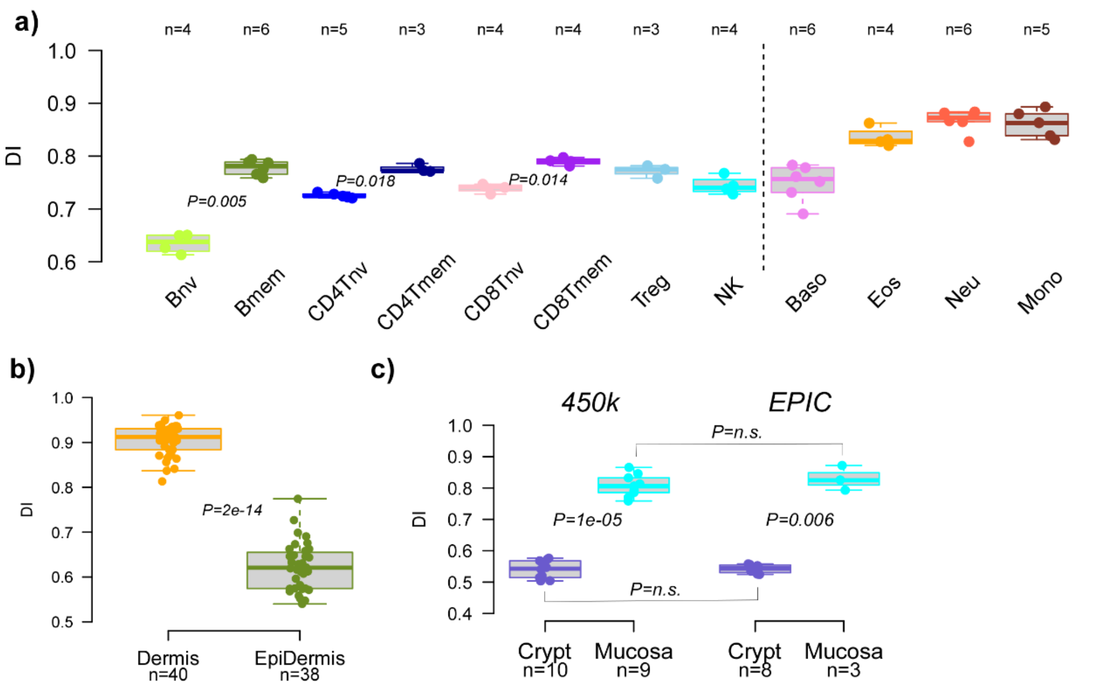


**SI fig.S4: DI varies within unipotent lineages and is robust to Illumina platform differences. a)** Boxplot displays the predicted DI values across 12 immune cell-types. EPIC DNAm data of sorted cells is from Salas et al Nat Commun 2022. Number of samples is indicated above plot. P-values derived from one-tailed Wilcoxon rank sum tests comparing naïve to memory counterparts in B-cells, CD4+ and CD8+ T-cells. **b)** Boxplots compare the predicted DI values in skin dermal (devoid of undifferentiated keratinocytes) vs epidermal (enriched for undifferentiated keratinocytes) samples, with the number of samples indicated below each plot and with the P-value derived from a one-tailed Wilcoxon rank sum test. **c)** Boxplots compare the predicted DI values in colon crypts (enriched for epithelial stem cells) vs colonic mucosa (enriched for differentiated progeny) samples, with the number of samples indicated below each box. P-values between crypt and mucosa samples derive from a one-tailed Wilcoxon rank sum test. P=n.s. not significant under a two-tailed Wilcoxon test.

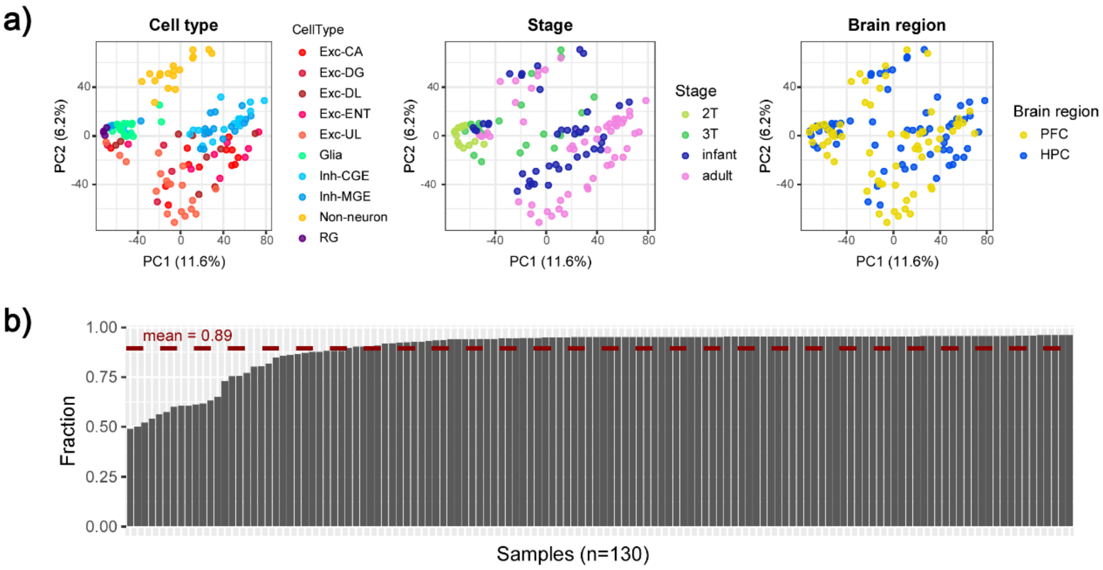


**SI fig.S5: Quality Control analysis of scnm3C-Seq3 brain data. a)** Scaterplot of PC1 vs PC2 from a PCA analysis on the 366,374 CpG-regions with > 90% coverage and 130 samples with > 50% coverage. Samples have been labeled by brain cell-type, developmental stage and brain region. **b)** Fraction of covered EpiDiff CpGs (n=1298) for each of the 130 samples, using a 2kb window centered at each EpiDiff CpG. Mean coverage is indicated by red dashed line.

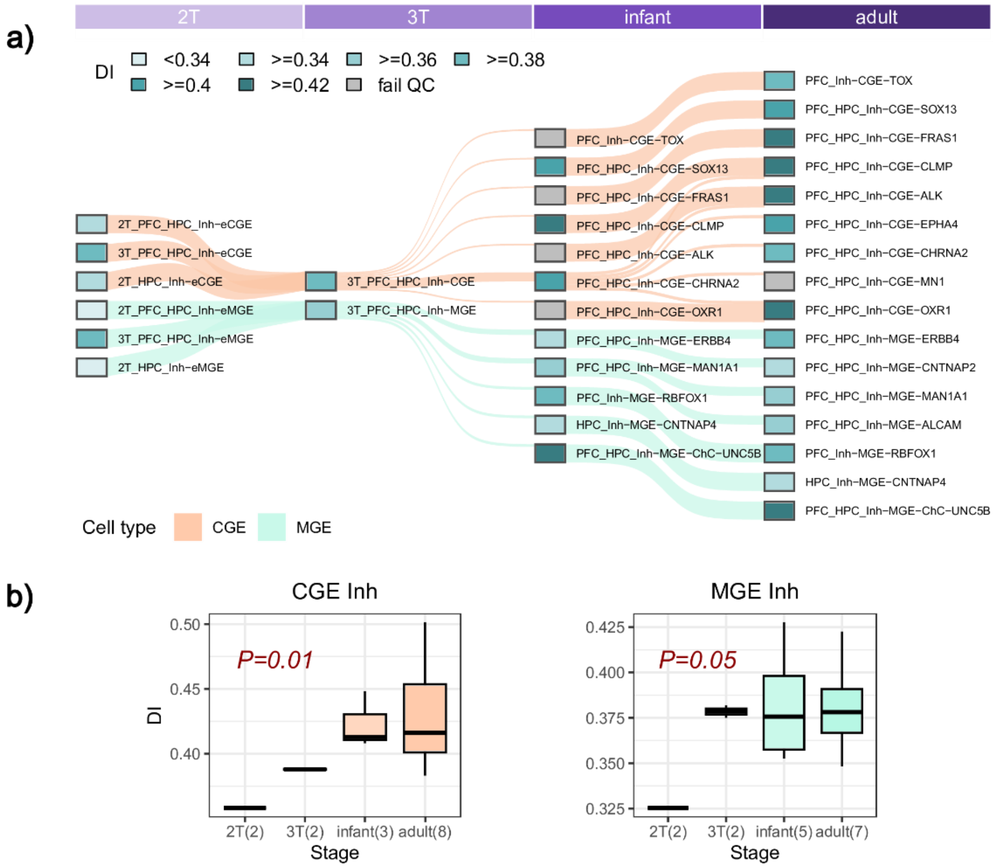


**SI fig.S6: EpiDiff predicts increased differentiation along inhibitory neuronal lineage in snm3C-Seq3 data. a)** Alluvial/Sankey diagram of the developmental lineage hierarchy of inhibitory neuronal cell-types during development of the human brain, with inhibitory neurons stratified by caudal or medial ganglionic eminence (CGE/MGE) regions. Developmental stages cover mid-gestation (2nd trimester-2T), late gestation (3rd trimester-3T), infants and adults. Each cell-type is labeled with a color according to their predicted DI-value, as indicated. Inh=inhibitory neuron. For full cell-type annotations, see Methods. **b)** Boxplots displaying the change in DI-values across developmental stages for the two inhibitory neuron lineages. P-values for CGE Inh and MGE inh lineages derive from a linear regression test.

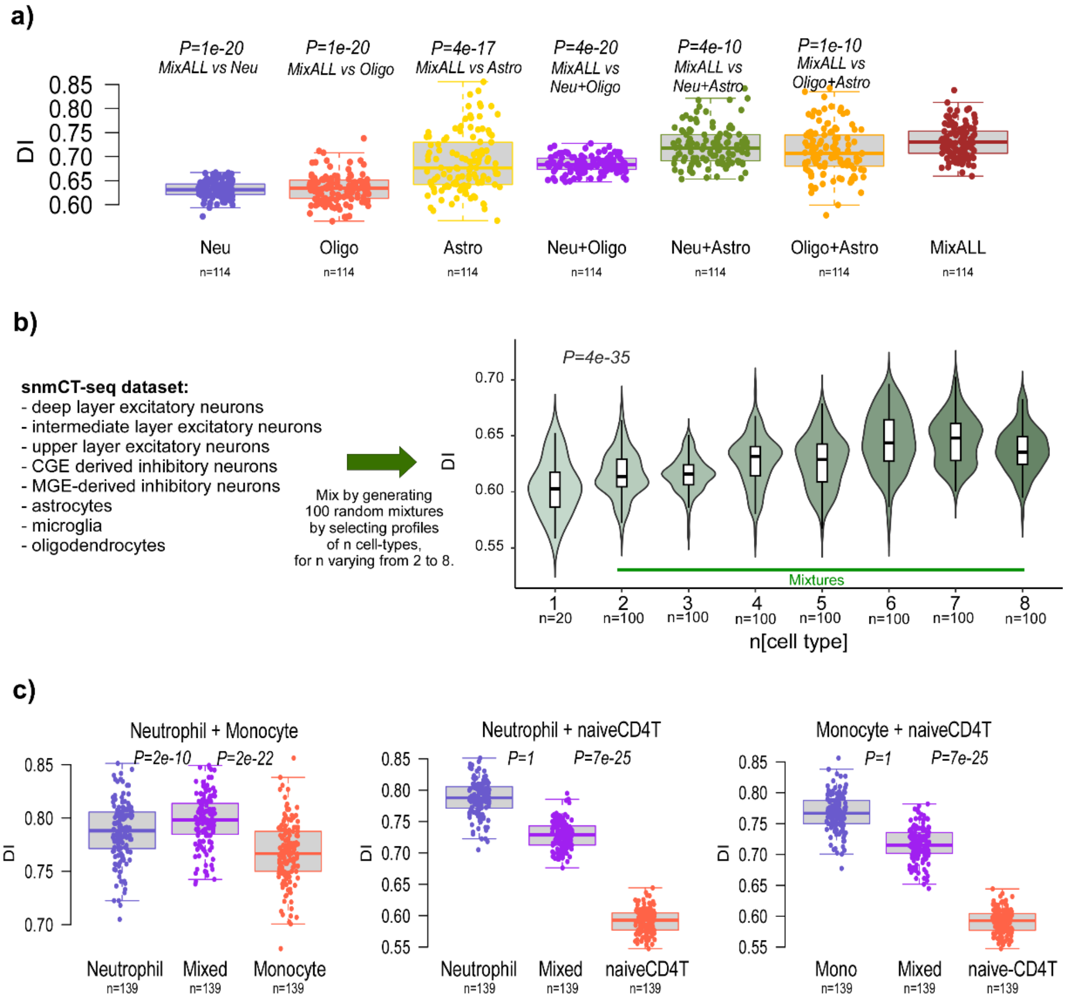


**SI fig.S7: Patterns of DI of cell-type mixtures. a)** DI of 114 neurons, 114 oligodendrocytes and 114 astrocytes, as well as their 2 and 3 cell-type mixtures, as indicated. There were 114 donors each contributing exactly 1 neuron, 1 oligodendrocyate and 1 astrocyte sample. Mixtures were generated using samples from the same donor to avoid confounding by genetics. P-values derived from a one-tailed paired Wilcoxon rank sum test, comparing the 3 cell-type mixture (MixALL) to the others. **b)** Violin plots display the DI values of 100 randomly selected mixtures at each value of “n” where n is the number of cell-types in the mixture. DNAm data is snmCT-Seq data, with a total of 8 cell-types as indicated on the left. When n=1, there are no mixtures and the violin describes the distribution over 20 samples, representing 8 distinct cell-types. P-value is from a linear regression of DI vs n. **c)** DI-values of sorted cell-types and their mixture, with DNAm data derived from BLUEPRINT. Each panel depicts a pair of cell-types with the “Mixed” sample describing a 50-50 mix of the underlying cell-types. There were 139 donors, each contributing 3 samples (neutrophils, naïve-CD4T and monocytes). When mixing we always mixed the sorted cell-type sample from the same donor. P-values in each panel compare the DI values of the mixed samples to the underlying sorted cell-types, using a one-tailed paired Wilcoxon rank sum test.

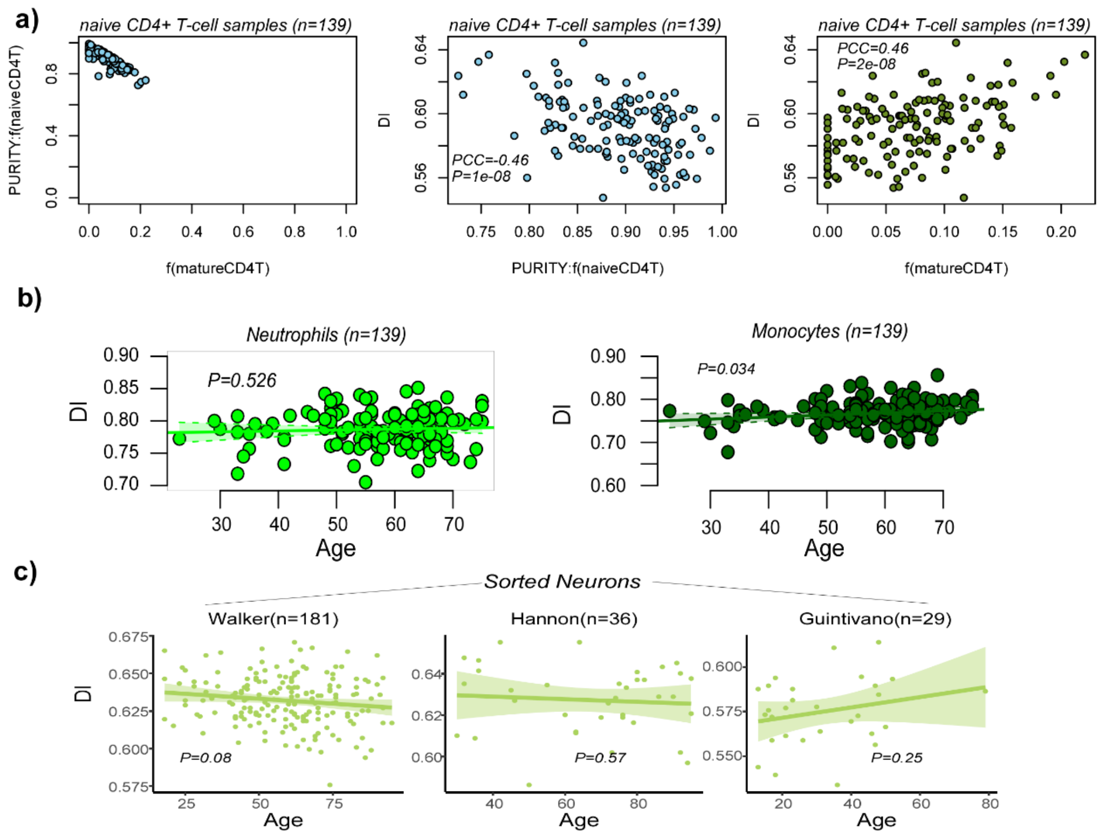


**SI fig.S8: DI associates with purity in naiveCD4T-cells and cell-types where DI does not correlate with age or only does so marginally (Illumina DNAm data). a)** Left panel: scatterplot of the estimated purity of the naïve CD4T cell samples against the estimated mature CD4T fraction in these same samples, with the estimated fractions obtained using EpiDISH with a 12 immune-cell type panel containing naïve and memory CD4T-cell subfractions. Middle and right panels: scatterplots of DI vs purity and estimated mature CD4T-cell fraction. Pearson Correlation Coefficient (PCC) and correlation-test P-value are given. **b)** Scatterplot of the DI from the Random Forest predictor against chronological age in the neutrophil and monocyte Illumina 450k dataset from BLUEPRINT. Fitted regression line and 95% confidence envelope is shown. P-value is from a linear regression. **c)** For three different Illumina DNAm datasets profiling sorted Neurons, we plot the predicted Differentiation Index (DI) vs chronological age. P-value from a two-tailed linear regression adjusted for sex is given. Number of samples in each set is given above panel. Fitted regression line and 95% confidence envelope is shown.

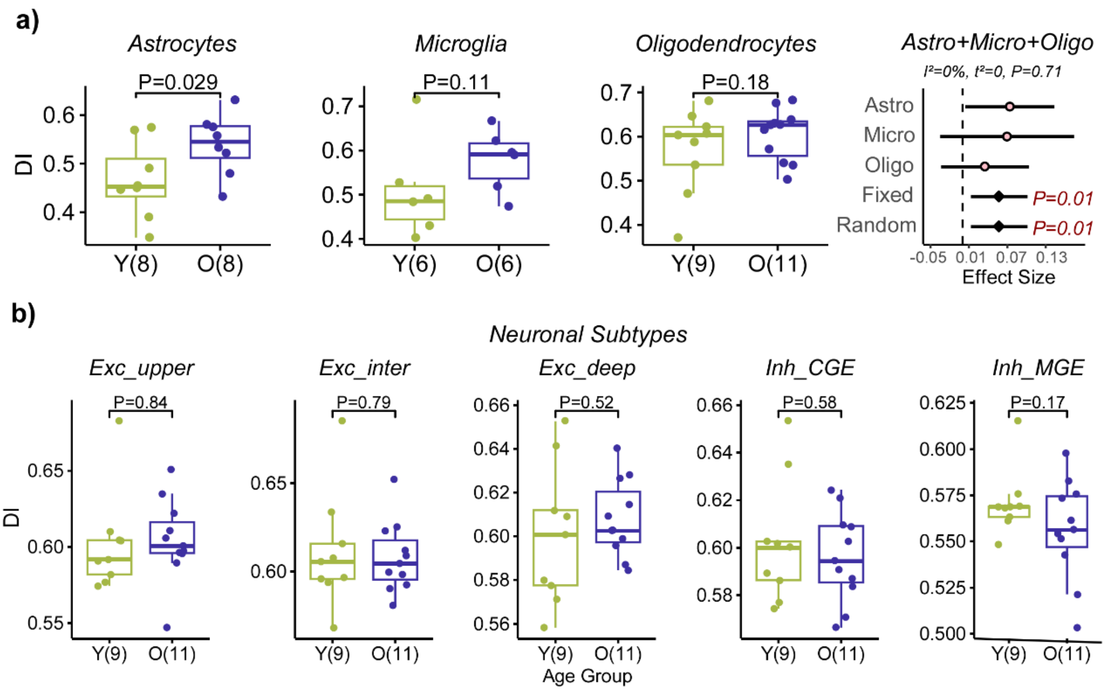


**SI fig.S9: Age-associated DI patterns in brain cell-types (snmCT-Seq data). a) Left panels:** Boxplots displaying predicted DI-values in young and old brain glia cell-types, where the DNAm data was snmCT-Seq. Number of independent young and old samples is indicated below each plot. P-value is from a one-tailed t-test. **Right panel:** Results of a fixed and random effect meta-analysis over the 3 separate glia subtypes, with effect size labeling the x-axis. For each datapoint in the Forest plot, the 95% confidence interval is given. P-values of the fixed and random effect models are given. Above the plot we display the heterogeneity indices and associated P-value of heterogeneity. **b)** As left panels in a) but for 5 neuronal subtypes as identified in a snmCT-Seq dataset, where the DI is being compared between 11 samples derived from old donors (O, ages>=70) and 9 samples derived from young donors (Y, ages<=30). P-value is from a two-tailed t-test.

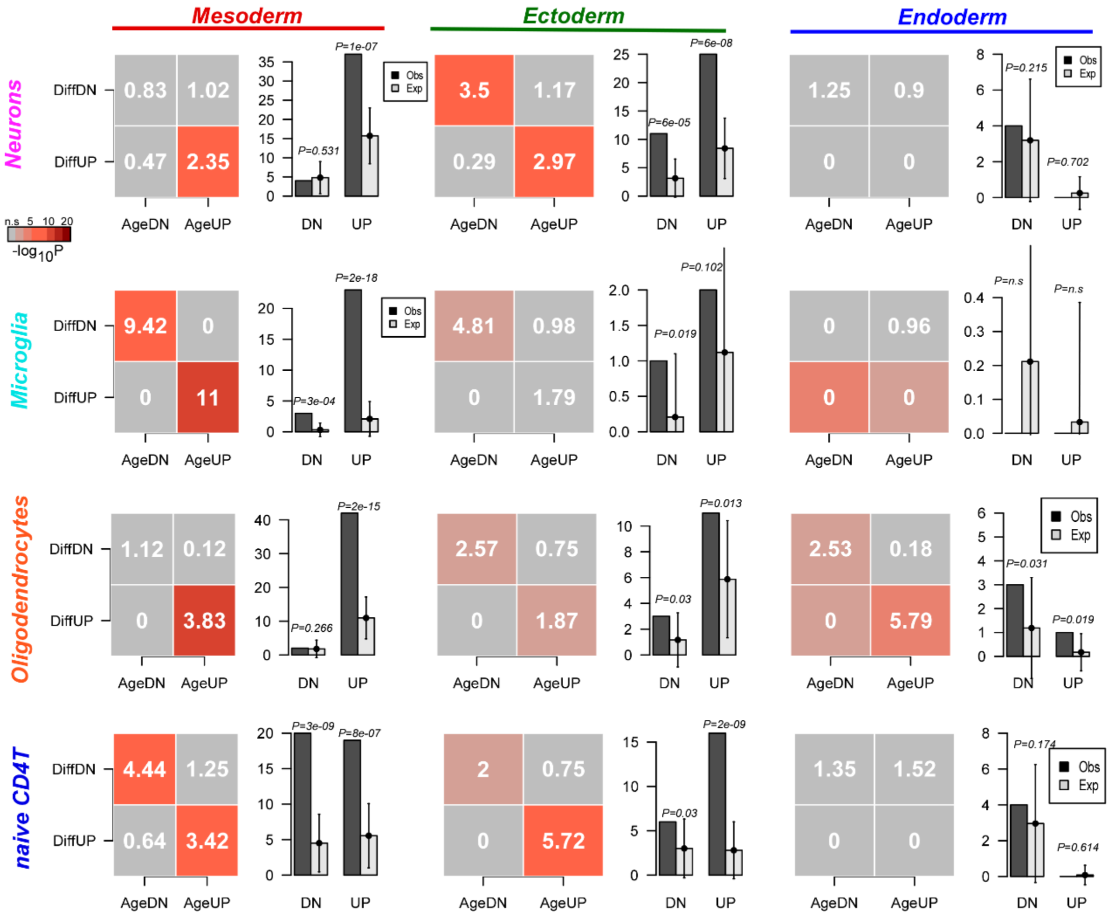


**SI fig.S10: Preferential mesoderm-drift in other cell-types.** For each germ-layer (mesoderm, ectoderm, endoderm) we display a heatmap of observed to expected ratios, one for each of four combinations of how germ-layer specific CpGs change with differentiation and with age in sorted neurons (n=181), microglia (n=12), oligodendrocytes (n=163) and naïve CD4+ T-cells (n=139). Significance for calling significant age-DMCs was determined at FDR<0.05. DiffUP/DiffDN: higher/lower methylation in fetal germ-layer compared to hESCs; AgeUP/AgeDN: age-associated hyper/hypo-methylation. The color in the heatmap indicates statistical significance level, as shown. Barplot on the right compares the observed (black) to expected (lightgrey) number of CpGs that exhibit hypermethylation with both differentiation and age (UP), or hypomethylation with both differentiation and age (DN). For the expected number, we also display the 95% confidence interval.

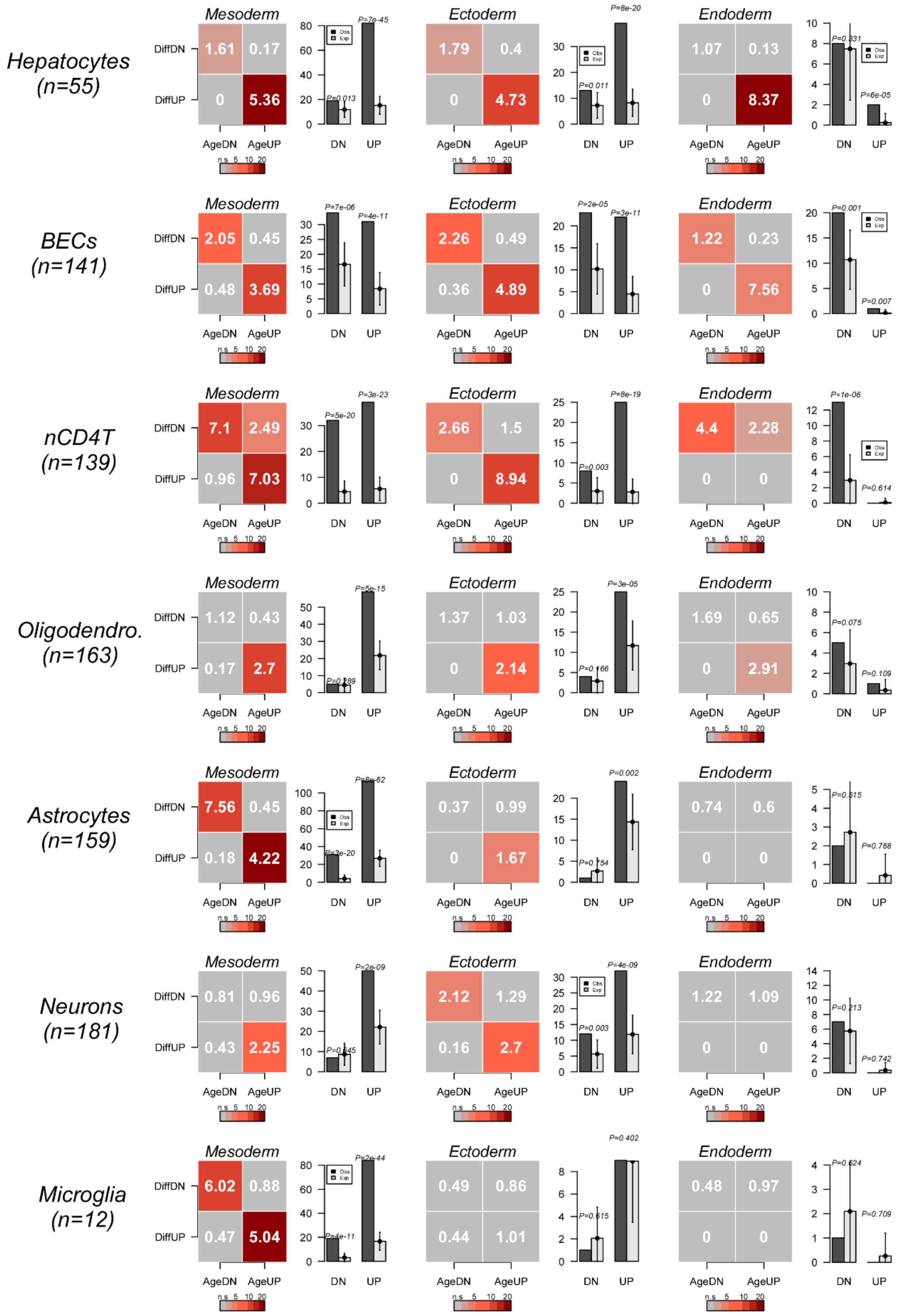


**SI fig.S11. Preferential mesenchymal-drift in 7 cell-types using nominal significance levels.** For each germ-layer (mesoderm, ectoderm, endoderm) we display a heatmap of observed to expected ratios, one for each of four combinations of how germ-layer specific CpGs change with differentiation and with age in a sorted cell-type, of which we display a total of seven. Number of samples of cell-type is indicated. Significance for calling significant age-DMCs was determined at a nominal $P < 0.05$ level. DiffUP/DiffDN: higher/lower methylation in fetal germ-layer compared to hESCs. AgeUP/AgeDN: age-associated hyper/hypo-methylation. The color in the heatmap indicates statistical significance level ($-\log_{10}P$). Barplot on the right compares the observed (black) to expected (lightgrey) number of CpGs that exhibit hypermethylation with both differentiation and age (UP), or hypomethylation with both differentiation and age (DN). For the expected number, we also display the 95% confidence interval.

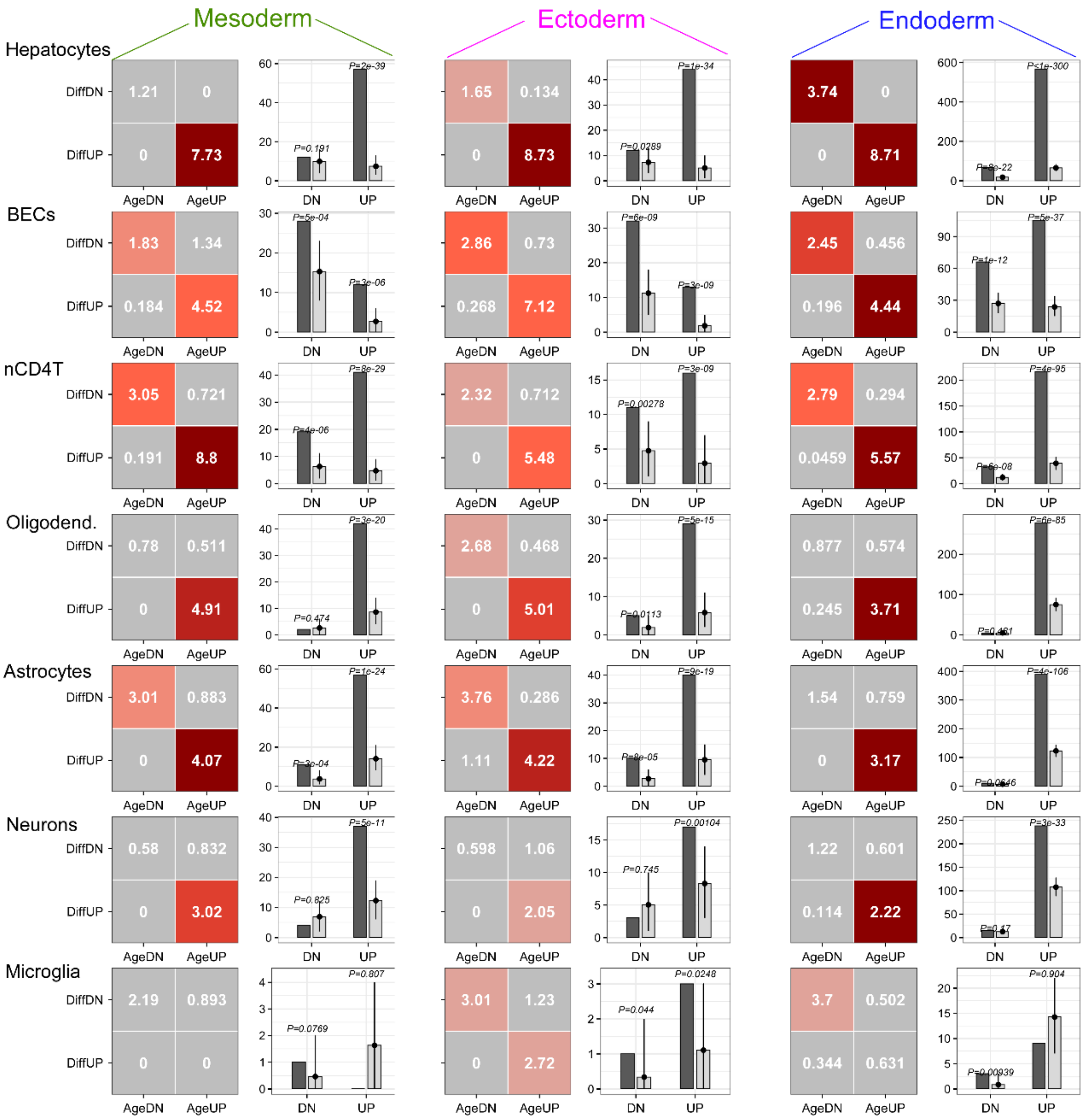


**SI fig.S12. Aligned developmental drift in 7 cell-types.** For each germ-layer (mesoderm, ectoderm, endoderm) we display a heatmap of observed to expected ratios, one for each of four combinations of how CpGs change with differentiation (adult vs fetal tissue within each germ-layer tissue originates from) and with age in a sorted cell-type, of which we display a total of seven. Number of samples of cell-type is indicated. Significance for calling significant age-DMCs was determined at FDR < 0.05 level. DiffUP/DiffDN: higher/lower methylation in adult vs fetal tissue within a given lineage. AgeUP/AgeDN: age-associated hyper/hypo-methylation. The color in the heatmap indicates statistical significance level (-$\log_{10}$P). Barplot on the right compares the observed (black) to expected (lightgrey) number of CpGs that exhibit hypermethylation with both differentiation and age (UP), or hypomethylation with both differentiation and age (DN). For the expected number, we also display the 95% confidence interval.

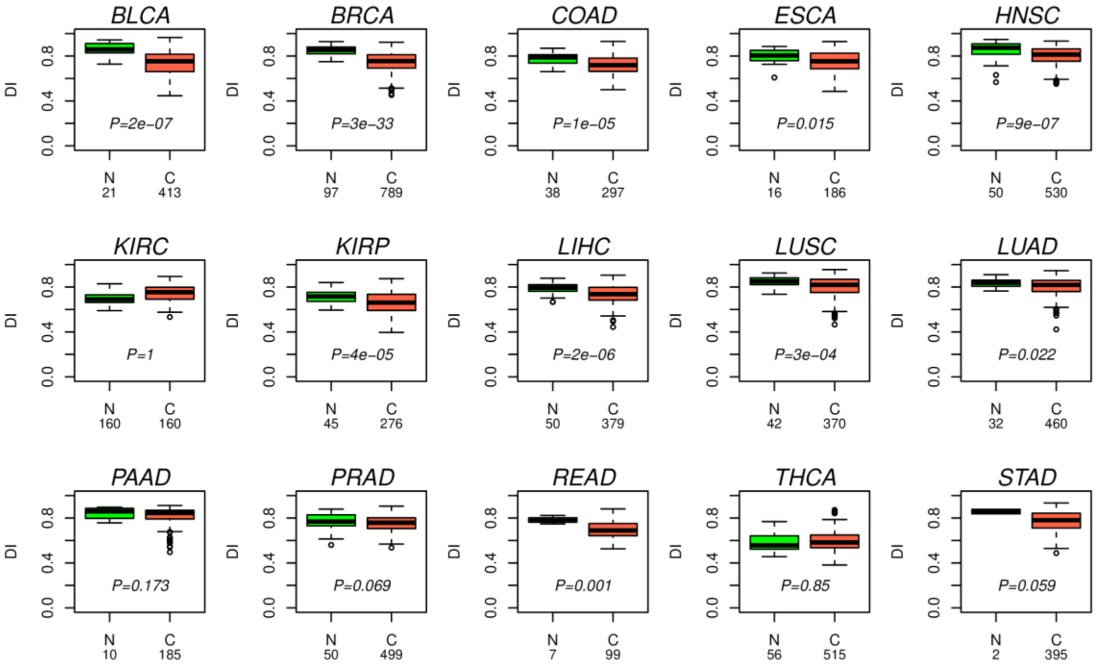


**SI fig.S13: The DI is reduced in TCGA cancer-types.** Boxplots compare the predicted differentiation index (DI) between normal-adjacent (N) and cancer-tissue (C) for 15 TCGA cancer-types. P-value is from a one-tailed Wilcoxon rank sum test. Number of samples in each category is given below x-axis.

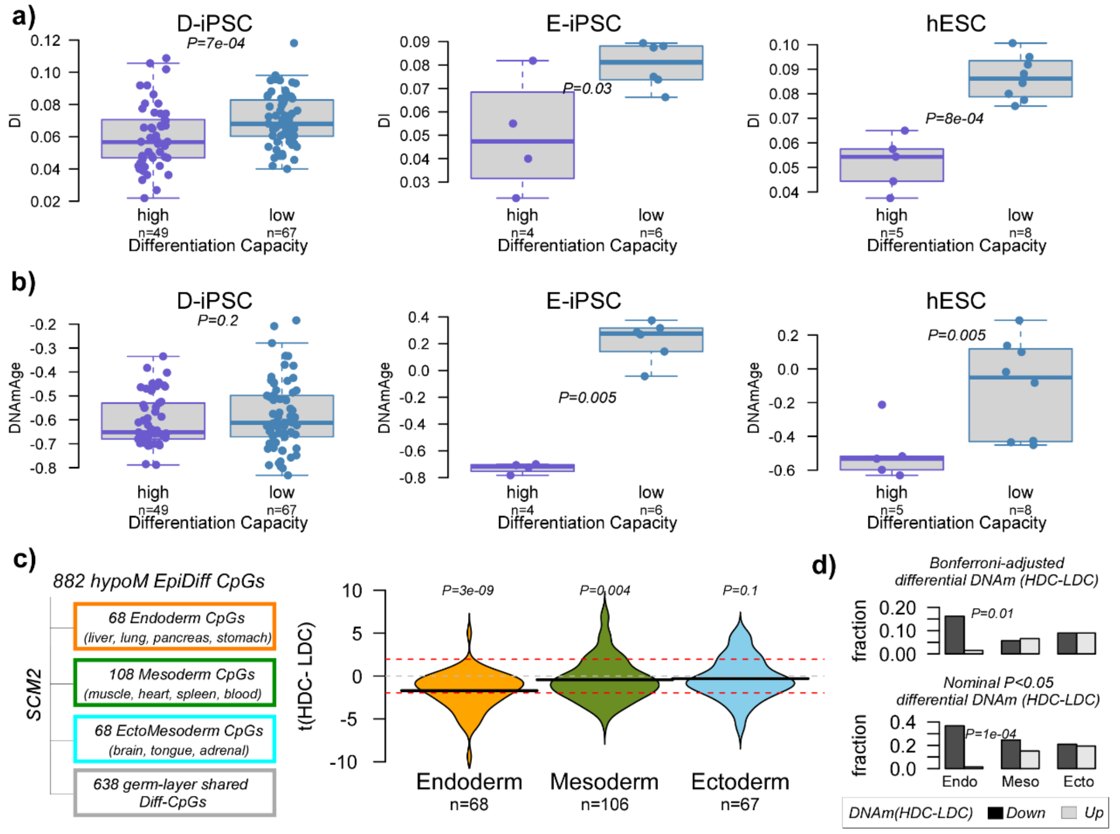


**SI fig.S14: DI correlates inversely with differentiation capacity. a)** Boxplots compare the predicted DI-values between iPSC or hESC cell-lines of low and high endodermal differentiation capacity (Butcher et al dataset). P-values are from one-tailed Wilcoxon tests. D-iPSC: iPSC derived from dermal fibroblasts, E-iPSC: iPSC derived from endothelial cells. **b)** As a) but for Horvath's DNAmAge. **c)** Definition of endoderm, mesoderm and ecto-mesoderm specific Diff-CpGs from the pool of 1298 EpiDiff-CpGs, as derived from EpiDiff's training dataset. These germ-layer specific DiffCpGs which become specifically hypomethylated in each respective germ-layer were then tested for differential DNAm in the Butcher et al dataset comparing high to low differentiation capacity (HDC-LDC) samples. Violins compare the t-statistics of differential DNAm (HDC-LDC) between the three sets of germ-layer specific CpGs. P-values are from a one-tailed Wilcoxon rank sum test to assess if the median is less than 0. Red dashed line are the lines of nominal P<0.05 significance. **d)** Barplots compare the fraction of germ-layer specific CpGs that display significant lower (down) or higher (up) DNAm levels in HDC compared to LDC samples, using a t-test, and using two different levels of statistical significance: Bonferroni adjusted P<0.05 and nominal P<0.05. P-values to assess difference in distribution derive from a one-tailed Fisher-test.
